# Accelerated Adaptation of SARS-CoV-2 Variants in Mice Lacking IFITM3 Preserves Distinct Tropism and Pathogenesis

**DOI:** 10.1101/2025.01.27.635150

**Authors:** Parker J. Denz, Jonathan L. Papa, Matthew I. McFadden, Prerana R. Rao, Jack Roettger, Adriana Forero, Jacob S. Yount

## Abstract

Here we investigated whether interferon induced transmembrane protein 3 (IFITM3), a key antiviral protein deficient in certain human populations, affects interspecies adaptation of SARS-CoV-2. We found that SARS-CoV-2 Beta and Omicron variants passaged through IFITM3-deficient versus wild type mice exhibit enhanced replication and pathogenesis in this new host species. Enhancements associated with amino acid substitutions in the viral genome, suggesting that IFITM3 limits accumulation of adaptive mutations. Mouse-adapted viruses enabled comparative studies of variants in mice. Beta caused lung dysfunction and altered cilia-associated gene programs, consistent with broad viral antigen distribution in lungs. Omicron, which shows low pathogenicity and upper respiratory tract preference in humans, replicated to high nasal titers while showing restrained spatial distribution in lungs and diminished lung inflammatory responses compared to Beta. Our findings demonstrate that IFITM3 deficiency accelerates coronavirus adaptation and reveal that intrinsic SARS-CoV-2 variant traits shape tropism, immunity, and pathogenesis across hosts.

**HIGHLIGHTS:**

- IFITM3 is a critical barrier to SARS-CoV-2 adaptation in new host species
- Mouse-adapted SARS-CoV-2 strains enable comparative pathology
- Omicron favors nose and large airways, leading to mild lung pathology
- Beta exhibits broad lung replication, driving severe inflammation and dysfunction

## INTRODUCTION

The emergence and rapid evolution of viruses in the human population highlights the continuous worldwide threat of respiratory virus spillover from animal reservoirs. Coronaviruses, such as SARS-CoV, MERS-CoV, and SARS-CoV-2, are believed to have emerged from animals, with subsequent adaptation enabling more efficient human-to-human transmission and severe disease^1–6^. Similar zoonotic transmission events have driven pandemics caused by influenza viruses, including the 1918 and 2009 H1N1 pandemics^7,8^. Despite these notable examples, pandemics are relatively rare even though humans are in constant contact with animals worldwide. This apparent paradox suggests the presence of significant barriers to zoonotic transmission and subsequent viral adaptation in humans. Understanding host and viral factors that facilitate or impede these processes will be critical for mitigating future pandemic risks.

Innate immune defenses, particularly those mediated by interferons, are a first line of protection against emergent viruses. The interferon-induced transmembrane protein 3 (IFITM3) is an antiviral protein that inhibits infection by numerous enveloped viruses by altering membrane properties of endosomes, thereby restricting virus-to-host membrane fusion^9–14^. Genetic deficiencies in IFITM3 have been associated with increased severity of both influenza virus and SARS-CoV-2 infections in humans^15–31^. In mouse models, IFITM3 deficiency similarly exacerbates disease severity for both viruses^15,32–35^. Additionally, in the context of influenza virus infection, loss of IFITM3 was shown to lower the minimum infectious dose threshold and to accelerate viral adaptation in mice^36^. These findings suggest that IFITM3 deficiencies could broadly decrease barriers to viral adaptation, but the effects of IFITM3 on the interspecies adaptation of other pandemic-relevant viruses remain unexplored.

In this regard, SARS-CoV-2 has undergone significant adaptation and evolution in humans since it emerged in 2019, resulting in viral variants with distinct properties. For example, the Beta variant has increased fusogenicity and pathogenicity compared to the ancestral SARS-CoV-2 strain, while Omicron variants are characterized by a preference for replication in the upper respiratory tract and reduced overall pathogenicity^37–42^. Notably, Omicron relies more heavily on endosomal entry than previous variants^37,38,40,41,43,44^, which may render it more sensitive to IFITM3-mediated restriction^9,32,37,45^, though this hypothesis has not been definitively tested *in vivo* since animal models have been limiting. Most SARS-CoV-2 variants only weakly infect young mice, and the gold-standard Syrian hamster model is poorly infected by Omicron variants, underscoring the need for new systems to better understand variant biology.

Here, we examined whether IFITM3 deficiency influences the interspecies adaptation of SARS-CoV-2 Beta and Omicron variants. By passaging these viruses through *Ifitm3^-/-^* mice, we demonstrate that the absence of IFITM3 facilitates the accumulation of species-specific adaptive mutations. We further compared the pathogenic properties of the new mouse-adapted variants generated in our study side-by-side with a commonly studied mouse-adapted virus derived from ancestral SARS-CoV-2 (MA10)^46,47^. Importantly, our mouse-adapted Beta and Omicron strains exhibit pathogenicity and tissue tropisms consistent with their unique profiles in humans, offering relevant and robust tools to study SARS-CoV-2 variant-specific pathology and immune responses in murine models. Global transcriptomic analysis of infected lungs revealed distinct inflammatory signatures, including strong alteration of cilia-associated genes and recruitment of neutrophils by Beta and MA10. Omicron infections, by contrast, elicited lower levels of lung transcriptional dysregulation and was restricted to the nose and large airways, explaining the milder pathology observed experimentally. Together, these findings provide critical insights into how IFITM3 influences SARS-CoV-2 adaptation and establish versatile new tools to investigate variant-specific host-pathogen interactions and immune responses *in vivo*.

## RESULTS

### IFITM3 deficiency facilitates adaptation of SARS-CoV-2 Beta variant to mice

To model an interspecies infection and adaptive evolution, we first compared mouse-adaptation of SARS-CoV-2 Beta variant upon lung-to-lung serial passaging after intranasal infection in WT versus IFITM3-deficient animals. The initial infection was administered at a dose of 10^5^ TCID50, followed by a 2-day replication period before collecting and homogenizing the lungs. The resulting lung homogenate fluid was then used to infect a subsequent mouse and this process was repeated for multiple serial passages (**Figure 1a**).

**Figure 1.**
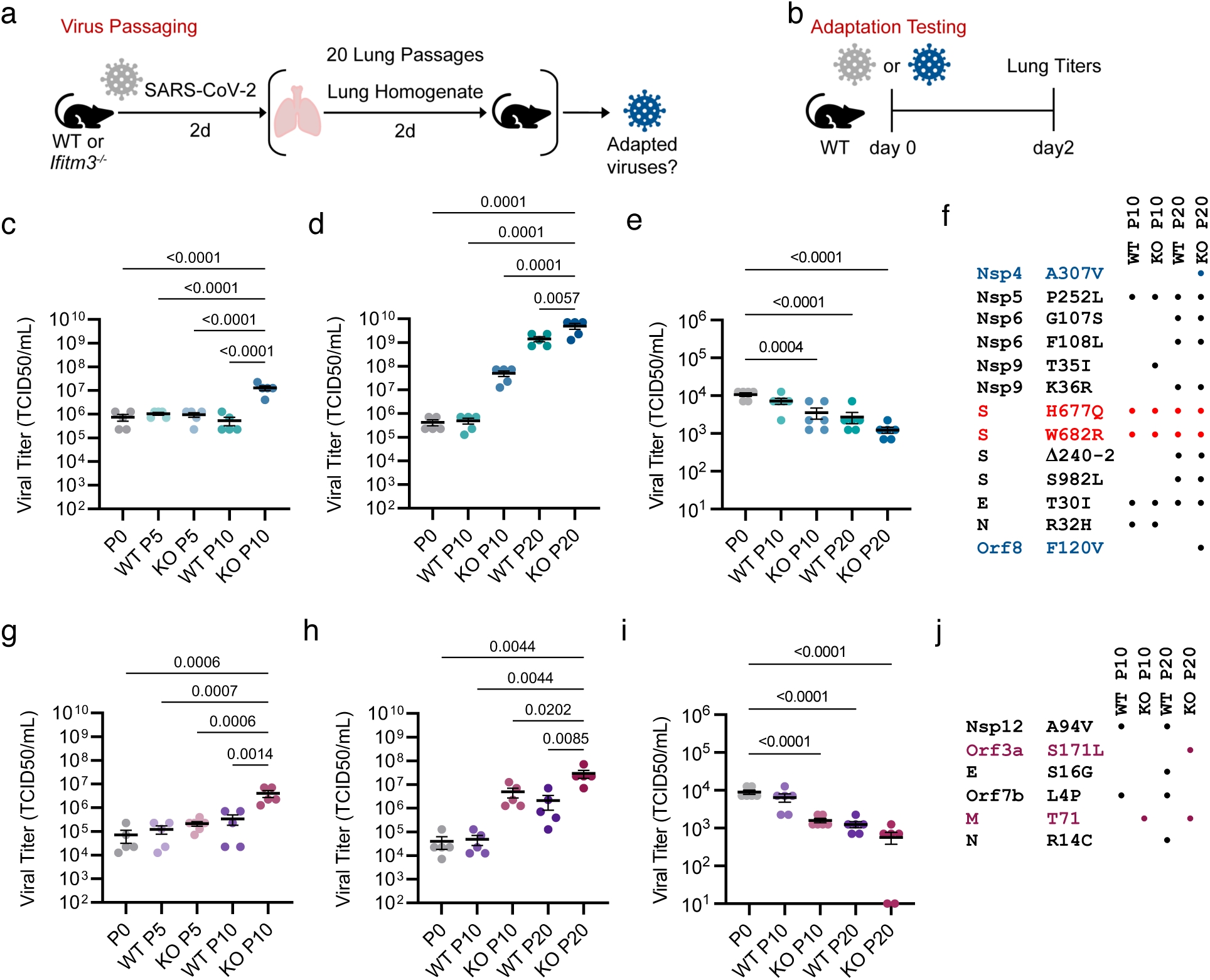
Passaging SARS-CoV-2 in the absence of IFITM3 enhances viral adaptation in mice. (**a**) Schematic of mouse passaging experiments. Initial intranasal infections were performed with 10^5^ TCID50 of the parental viruses. (**b**) Schematic of WT mouse challenge with parental SARS-CoV-2 Beta, Omicron, or passaged viruses. (**c, d, g, h**) Groups of WT mice (n=5 per group) were challenged with 10^5^ TCID50 of the SARS-CoV-2 Beta (**c, d**) or Omicron (**g, h**) strain passaged 5, 10, or 20 times through WT or *Ifitm3^-/-^* mice and compared to the respective parent virus (passage 0). (**e,i**) A549-ACE2 cells were infected with the indicated viruses at an MOI of 1 and incubated for 48 hours to allow for multi-cycle replication. The supernatants were collected to determine viral titer. Titer data is representative of 2 independent experiments each performed in triplicate (n=6). (**f, j**) Table of amino acid substitutions found in the consensus sequence of the Beta (mutations specific to KO passage 20 are shown in blue and furin cleavage site reversions are shown in red) (**f**) or Omicron (KO passage 20-specific mutations are shown in purple) (**j**) viruses after passaging 10 times or 20 times through WT of *Ifitm3^-/-^* mice.

We propagated virus passages 5 and 10 from both WT and *Ifitm3*^-/-^ mice, as well as the parental virus stock (passage 0), in Vero-TMPRSS2 cells, and used these expanded stocks to challenge groups of WT mice with equal virus doses. The mice were sacrificed on day 2 post infection to measure viral titers (**Figure 1b**). We found that mice challenged with KO passage 10 were the only group with a statistically significant increase in lung viral titers compared to the parent virus infection (**Figure 1c**). These results showed that 10 virus passages through IFITM3-deficient mice resulted in detectable viral adaptation to mice and indicate that IFITM3 is a significant bottleneck for interspecies adaptation.

To determine whether additional passages would enhance adaptation or allow adaptation in WT mice, we continued our lung-to-lung passaging an additional 10 times, resulting in 20 total passages through WT or *Ifitm3^-/-^* mice. Groups of WT mice were then infected with 10^5^ TCID50 of WT or KO passages 10 and 20 in comparison to the parental Beta virus. KO passage 20 replicated to the highest levels compared to all other stocks, indicating that the greatest adaptation continued to be seen for *Ifitm3^-/-^*-passaged virus (**Figure 1d**). However, WT passage 20 also gained an increase in viral titers compared to the parent virus, suggesting that significant viral adaptation had also occurred but was delayed by the expression of IFITM3 (**Figure 1d**). We also tested the passaged virus stocks for replication fitness in A549 human lung cells stably expressing human ACE2. Replication in the human cells was significantly reduced for virus passaged 20 times through either WT or *Ifitm3^-/-^* mice, indicating that mouse adaptation led to decreased viral fitness in human cells (**Figure 1e**). These results identify a critical role of IFITM3 in the restriction of interspecies adaptation of SARS-CoV-2 and highlight tradeoffs of enhanced replication in mice with respect to replicative fitness in human cells.

### IFITM3 deficiency facilitates the accumulation of adaptative mutations in the SARS-CoV-2 Beta variant

We next extracted viral RNA and sequenced the genomes of WT and KO passages 10 and 20 in comparison to the parental Beta strain. Sequencing confirmed the identity of the Beta parent strain, which is characterized by many mutations across the genome as compared to the WA1 strain, including the N501Y mutation in the Spike protein that allows interaction with murine ACE2^48^ (**Supplementary Figure 1a**). Our stock of parent Beta virus also contained 2 tissue culture adaptations (>90% prevalence) in or near the furin cleavage site of the S protein that are known to arise when propagating the virus in Vero cells (**Supplementary Figure 1a**)^49–51^.

We observed clear changes within the consensus sequences (>50% prevalence) of the passaged viruses. Several mutations were shared between both WT passage 10 and KO passage 10, including Nsp5 P252L, E T30I, N R32H and 2 Spike changes that fully reverted the tissue culture-associated mutations at the furin cleavage site, H677Q and W682R (**Figure 1f**, **Supplementary Figure 1b, c**). Importantly, the presence of an intact furin cleavage site was not sufficient alone to increase virus replication since this site was fully restored (no mutant sequences detected) by 10 passages in either WT or KO mice while only KO passage 10 showed a significant increase in virus replication (**Figure 1c**). KO passage 10 had a single unique mutation within nsp9 (T35I), a protein critical in the formation of the viral replication complex^52^, which associates with its enhanced replication (**Figure 1f**, **Supplementary Figure 1c**).

Analyzing sequences of the 20^th^ passage through WT or *Ifitm3^-/-^*mice, we again saw several shared mutations, including 2 nsp6 mutations (G107S, F108L), a new nsp9 mutation (K36R) that seems to have replaced the one present in KO passage 10, a three-residue deletion in Spike (amino acids 240-242), and a Spike S982L mutation (**Figure 1f**, **Supplementary Figure 1d, e**). The increased replication levels observed in mice challenged with the KO passage 20 Beta strain were associated with 2 unique mutations, Nsp4 A307V and ORF8 F120V (**Figure 1f**, **Supplementary Figure 1e**). These data indicate that while mutations associated with adaptation were selected in both WT and *Ifitm3^-/-^* mice, this process was enhanced in the absence of IFITM3.

### IFITM3 deficiency facilitates adaptation of SARS-CoV-2 Omicron BA.4 variant to mice

Omicron variants preferentially enter cells via endocytosis, which has been proposed to make them more susceptible to IFITM3 restriction in late endosomes compared to the earlier variant strains, such as Beta, which fuse preferentially at the cell surface^37,38,43^. We thus examined whether Omicron BA.4 virus showed distinct replication in WT versus *Ifitm3^-/-^* mice. We observed a 2-log increase in viral titers in the lungs of *Ifitm3^-/-^* mice versus WT mice, demonstrating that Omicron is highly sensitive to IFITM3 restriction *in vivo* (**Supplementary Figure 2a**). We also observed increased viral titers in the noses of the infected *Ifitm3^-/-^*mice (**Supplementary Figure 2b**). Corresponding with viral loads, inflammatory cytokine levels were similarly elevated in the KO lung samples (**Supplementary Figure 2c**). The low level of replication and cytokine induction in WT mice suggested that the Omicron strain retains significant potential for mouse adaptation.

To explore this further, we performed a similar set of passaging experiments through WT and *Ifitm3^-/-^* mice using the BA.4 Omicron variant. When challenging groups of WT mice with equal doses of passages 5 and 10, we saw that the Omicron virus passaged 10 times through *Ifitm3^-/-^* mice replicated to the highest levels in mouse lungs, indicating significant mouse adaptation (**Figure 1g**). We continued our lung-to-lung passaging an additional 10 times, totaling 20 passages through WT or *Ifitm3^-/-^* mice, to determine whether a virus with greater adaptation could be obtained. Testing WT and KO passages 10 and 20, we observed that KO passage 20 had the highest levels of virus present in the lungs, though like Beta passages, the WT passage 20 Omicron virus showed replication levels that fell between KO passages 10 and 20, indicating that substantial adaptation had also occurred via WT mouse passaging (**Figure 1h**). In human A549-ACE2 lung cells, viruses passaged through either WT or *Ifitm3^-/-^* mice showed reduced levels of replication compared to the parent virus with 2 out of 6 KO passage 20 replicate samples showing levels of replication below the detection limit of the TCID50 assay (**Figure 1i**). These experiments corroborate the conclusion that IFITM3 deficiency facilitates species-specific adaptation of distinct SARS-CoV-2 strains in mice, regardless of their preferred cell entry mechanisms.

### IFITM3 deficiency facilitates accumulation of adaptative mutations in the SARS-CoV-2 Omicron BA.4 Variant

We next extracted viral RNA and sequenced the genomes of the Omicron virus passaged 10 and 20 times through WT or *Ifitm3^-/-^* mice in comparison to the parental Omicron virus. The Omicron lineage of SARS-CoV-2 has more mutations than any previous variant, with at least 20 mutations within the Spike protein alone, and our sequencing confirmed the identity of the parent virus (**Figure 1j, Supplementary Figure 3a**)^39^. WT passage 10 had two unique mutations, one in nsp12 (A95V) and one in ORF7b (L4P), but these were not associated with an increase in viral replication (**Figure 1j, Supplementary Figure 3b**). Interestingly, the increase in viral replication by KO passage 10 is linked to a single unique amino acid mutation within the M gene, T7I (**Figure 1j, Supplementary Figure 3c**). WT passage 20 retained the 2 mutations that arose in passage 10 and gained mutations in the N protein (R14C) and E protein (S16G) (**Figure 1j, Supplementary Figure 3d**). KO passage 20 retained the T7I mutation in the M protein and acquired a single new mutation in ORF3a (S171L) (**Figure 1j, Supplementary Figure 3e**). These Omicron passaging experiments, coupled with data from our Beta variant experiments, demonstrate that deficiency in IFITM3 allows for rapid adaptation of SARS-CoV-2 to a new host species via the accumulation of specific mutations.

### Enhanced inflammatory responses across mouse-adapted SARS-CoV-2 variants

Having established that IFITM3 deficiency accelerates the adaptation of SARS-CoV-2 Beta and Omicron variants to mice, we next sought to assess the functional consequences of these adaptations on host pathology and immune responses. We measured weight loss as an indicator of overall disease severity and analyzed inflammatory cytokine levels in the lungs to determine the impact of passaged viruses on host immune activation.

We first examined whether early signs of adaptation could be detected at passage 5. For the Beta variant, neither WT nor KO passage 5 viruses induced detectable weight loss, but the KO passage 5 virus elicited a subtle yet significant increase in IL-6, suggesting initial adaptive changes (**Supplementary Figure 4a,b**). In contrast, weight loss and cytokine induction became readily apparent for later passages. While WT passage 10 induced minimal weight loss, KO passage 10 induced moderate weight loss (**Figure 2a**). KO passage 20 induced the greatest amount of weight loss and WT passage 20 showed an intermediate weight loss phenotype (**Figure 2a**). These weight loss results generally correlated with viral titers shown in **Figure 1d**. Largely mirroring the weight loss and viral titers, levels of the inflammatory cytokines IFNβ, IL-1β, TNF, and IL-6 were highest in the lungs of mice infected with KO passage 20, though KO passage 10 and WT passage 20 also induced robust increases compared to the parent virus (**Figure 2b**). Since the pro-inflammatory cytokines we measured are known to drive inappetence and weight loss^32,33,53–56^, the more robust cytokine responses observed at passages 10 and 20 may underlie the more pronounced weight loss phenotypes induced by the later passages.

**Figure 2.**
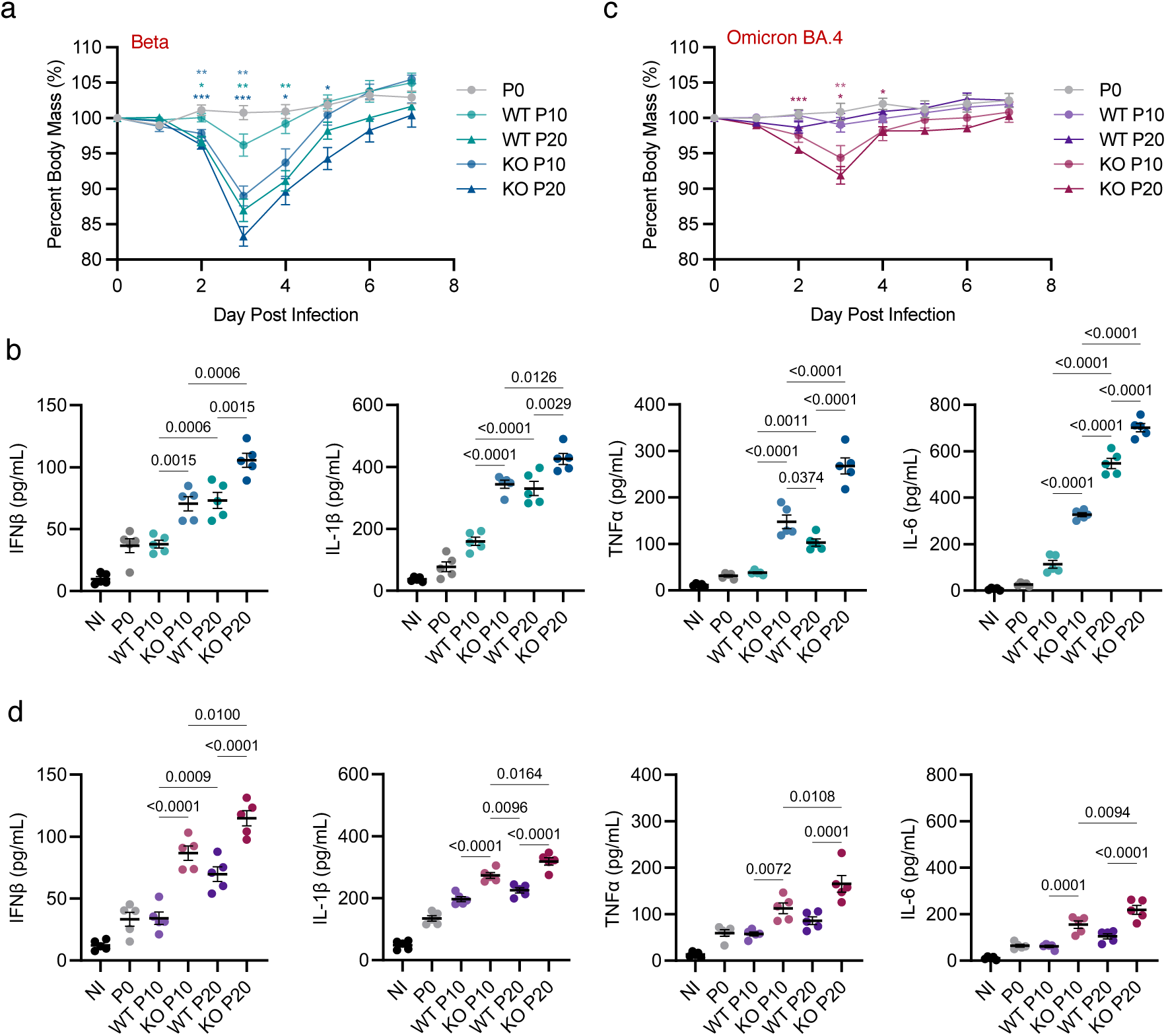
SARS-CoV-2 variants passaged in the absence of IFITM3 show enhanced pathogenicity in mice. Groups of WT mice (n=5 per group) were challenged with 10^5^ TCID50 of the SARS-CoV-2 Beta (**a, b**) or Omicron (**c, d**) strain passaged 10 or 20 times through WT or *Ifitm3^-/-^*mice and compared to the parent virus (passage 0). (**a, c**) Weight loss during challenge by passaged SARS-CoV-2 viruses. Error bars represent SEM, comparisons were made using the Mann-Whitney test. (only comparisons to P0 are shown; p values: **** <0.0001, *** <0.0005, ** <0.005, *<0.05). (**b, d**) ELISA quantification of IFNβ, IL-1β, TNFα, and IL-6 in lung homogenates collected at day 2 post infection. Error bars represent SEM and comparisons were analyzed by ANOVA followed by Tukey’s multiple comparisons test. (**a, c**) Dots represent averages of individual mice (n=5 per group). (**b, d**) Each dot represents an individual mouse.

We observed similar trends for the Omicron variant. Neither the parental Omicron virus nor WT passages 5 or 10 induced appreciable weight loss or cytokine increases (**Supplementary Figure 4c,d, Figure 2c**). However, KO passages 10 and 20 both induced moderate weight loss (**Figure 2c**) as well as the highest levels of lung IFNβ, IL-1β, TNF, and IL-6 (**Figure 2d**). WT passage 20 also induced a moderate increase in these cytokines relative to the parental virus (**Figure 2d**), though it did not induce weight loss (**Figure 2c**). These results show that serial passage in IFITM3-deficient mice drives viral adaptation that results in heightened inflammatory responses and morbidity for both Beta and Omicron variants. Moreover, these findings indicate that modest increases in pro-inflammatory cytokines are not sufficient to induce overt weight loss and that surpassing a threshold of cytokine induction may be required for more pronounced disease manifestations.

### Comparing replication, tropism, and pathology of adapted SARS-CoV-2 variants in mice

The viruses we have generated now enable direct comparative biology of SARS-CoV-2 variants in mice, providing a unique opportunity to study the replication, tropism, and pathology of Beta and Omicron strains alongside the well-characterized MA10 mouse-adapted strain derived from the earliest SARS-CoV-2 isolate^46,47^. To examine variant-specific characteristics, we compared the most highly adapted Beta and Omicron strains (WT and KO passage 20) to MA10 in young C57BL/6 mice. We found that mice infected with MA10 lost the least amount of weight among the three infections, while Omicron induced moderate weight loss and Beta induced a more severe drop in weight with a longer time to recovery (**Figure 3a**). Lung viral titers at day 2 were highest for Beta-infected mice, followed by Omicron and MA10 (**Figure 3b**). We also examined nasal titers given the reported preference of Omicron for the upper respiratory tract^37,39,40,42,44^ and indeed observed the highest average nose titers for Omicron infection (**Figure 3c**). Beta titers were also relatively high in the nose in comparison to MA10 titers that were more than 1 log lower (**Figure 3c**). We also examined heart titers for the three viruses since acute and long-term cardiac complications are among the most prevalent extrapulmonary manifestations of SARS-CoV-2 infections. MA10 was not detected in the heart, consistent with past reports from us and others (**Figure 3d**)^32,46,47^. However, Beta and Omicron were both present at detectable levels in heart tissue, suggesting dissemination and cardiac replication of these viral variants (**Figure 3d**).

**Figure 3.**
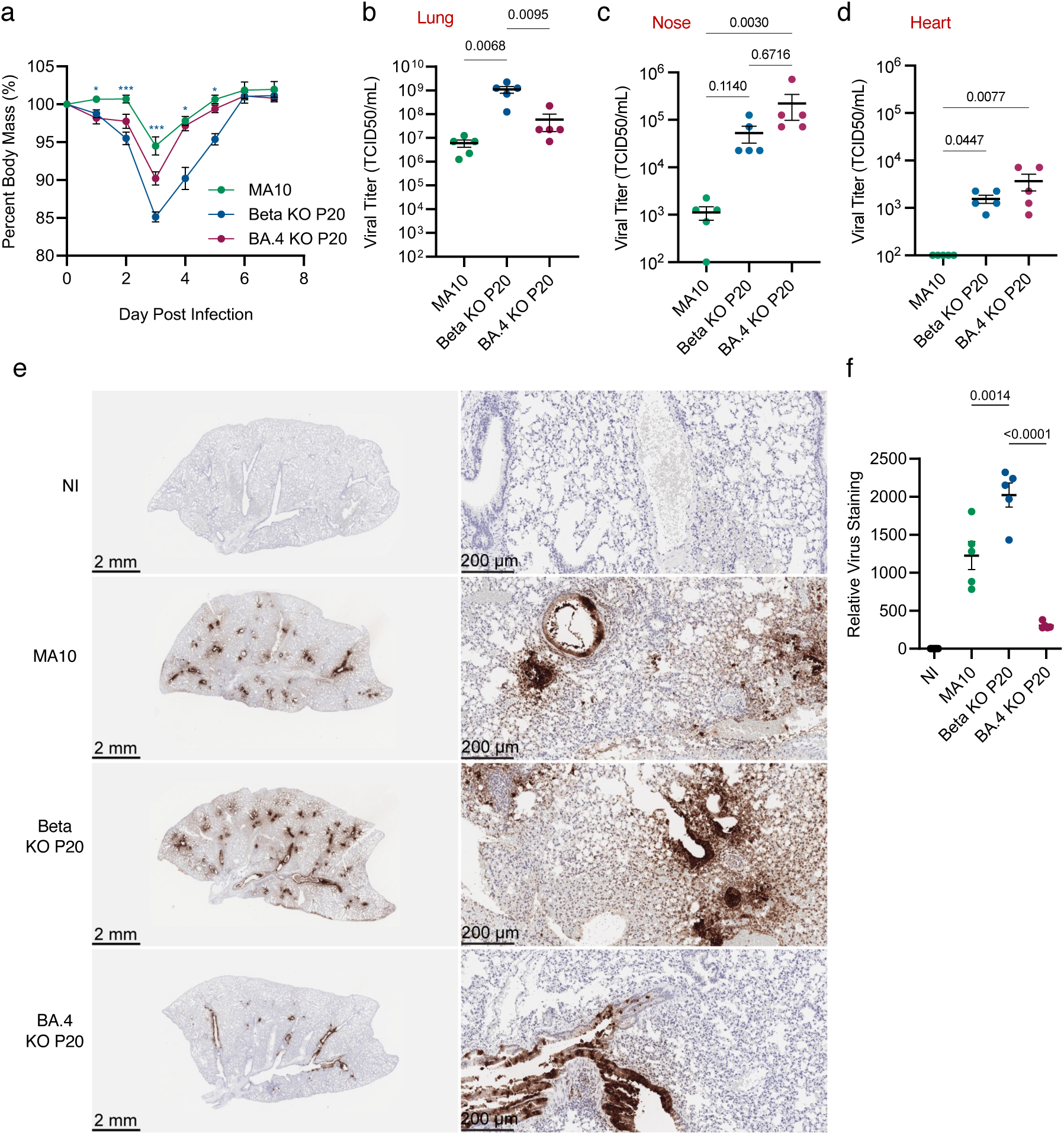
Comparative analysis of mouse-adapted Beta, Omicron, and MA10 strains. Groups of WT mice (n=5 per group) were challenged with 10^5^ TCID50 of the SARS-CoV-2 BA.4 strain or Beta strain passaged 20 times through *Ifitm3^-/-^*mice and compared to infection by an equal dose of MA10. (**a**) Weight loss during challenge by passaged SARS-CoV-2 viruses. Error bars represent SEM, comparisons were made using the Mann-Whitney test (p values: **** <0.0001, *** <0.0005, ** <0.005, *<0.05). Viral titers from lung (**b**), nose (**c**), or heart (**d**) homogenates collected at day 2 post infection. (**e**) Representative images of infected lungs at day 2 post infection stained with IHC staining using an anti-SARS-CoV-2 Nucleocapsid antibody. Whole lung images are shown on the left (scale bars represent 2 mm) and 10x zoomed images are shown on the right (scale bars represent 200 μm). (**f**) ImageJ was utilized to quantify the lung section staining for entire lung sections of multiple infected animals at day 2 post infection. (**a**) Dots represent averages of individual mice (n=5 per group). (**b-d, f**) each dot represents an individual mouse, *n* = 5, error bars indicate SEM, significance determined by one-way ANOVA. Numbers above the graph represent p values.

Replication of Omicron to relatively robust levels in the lungs was somewhat unexpected, given previous reports describing its limited lung involvement in humans^37–41,44^. To explore this further, we examined viral antigen staining at day 2 post infection in lung tissue and observed distinct patterns across MA10, Beta, and Omicron infection conditions. MA10-infected lungs exhibited discrete, focal clusters of staining predominantly surrounding airways and vascular structures (**Figure 3e**). Beta-infected sections displayed the most extensive and intense viral antigen staining, involving both alveolar and bronchial regions (**Figure 3e**). In contrast, Omicron-infected samples showed moderate staining, primarily concentrated around bronchi and bronchioles, with minimal involvement of alveolar spaces or areas outside larger airways (**Figure 3e**). Unbiased quantification of lung sections from multiple mice confirmed that Omicron-infected lungs showed the least viral antigen staining followed by MA10 and Beta (**Figure 3f**). These images highlight two key concepts: (1) Omicron-infected lung sections contained fewer infected cells compared to other variants, consistent with a reduced tropism for lung tissue. However, given the robust infectious viral titers detected in these lungs, this suggests a higher viral output per infected cell, aligning with recent studies^41^. (2) We additionally observed that densely packed antigen-positive cells, likely representing dead or dying cells, were obstructing airways in MA10- and Beta-infected samples (**Figure 3e**). These observations suggest that MA10 and the adapted Beta variant likely cause greater airway dysfunction in mice than adapted Omicron.

To test this, we measured lung function by whole body plethysmography during a time course of infection. We saw that the PenH, an indicator of airway resistance, significantly increased for both MA10 and Beta KO passage 20, with Beta inducing the highest PenH readings (**Figure 4a**). In contrast, Omicron infection induced a nearly imperceptible change in PenH, indicating preserved airway function despite its replication in lung tissue (**Figure 4a**).

**Figure 4.**
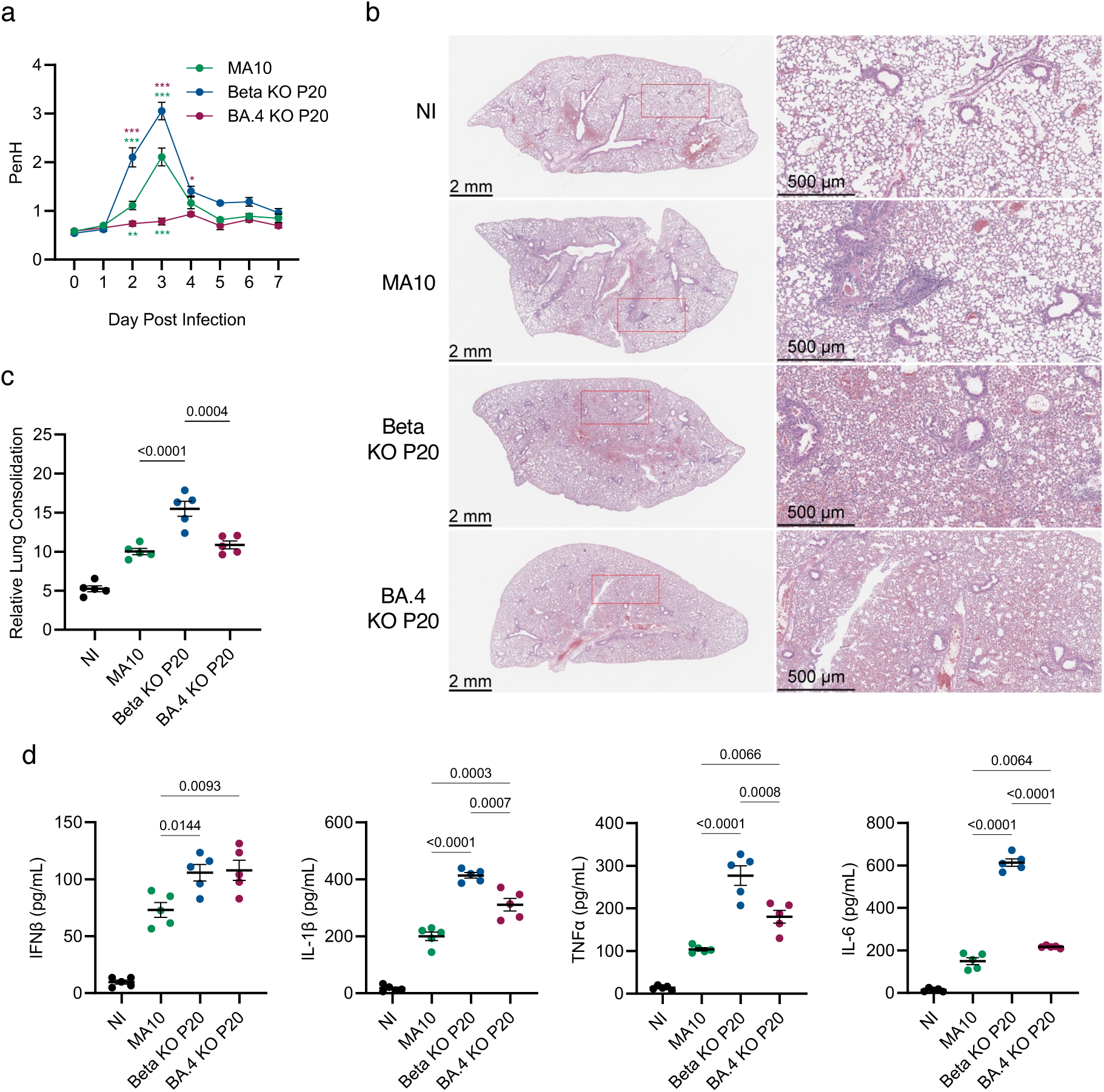
Comparing lung pathogenesis and inflammation between mouse-adapted Beta and Omicron strains to MA10. Groups of WT mice (n=5 per group) were challenged with 10^5^ TCID50 of the SARS-CoV-2 BA.4 strain or Beta strain passaged 20 times through *Ifitm3^-/-^* mice and compared to infection by an equal dose of MA10. (**a**) Daily whole-body plethysmography enhanced pause (PenH) measurements. Dots represent averages of individual mice (n=5 per group) (p values: **** <0.0001, *** <0.0005, ** <0.005, *<0.05). (**b**) Representative histology images of infected lungs at day 7 post infection. Whole lung images are shown on the left (scale bars represent 2 mm) and 4x zoomed images are shown on the right (scale bars represent 500 μm). The red box present in the whole lung images corresponds to its matched zoomed in image. (**c**) Quantification of consolidated tissue versus open airspace in H&E staining for entire lung sections of multiple animals infected with Beta KO Passage 20, BA.4 KO Passage 20 or MA10 at day 7 post infection. (**d**) ELISA quantification of IFNβ, IL-1β, TNFα, and IL-6 in lung homogenates collected at day 2 post infection. Error bars represent SEM and comparisons were analyzed by ANOVA followed by Tukey’s multiple comparisons test. (**c, d**) Each dot represents an individual mouse. Numbers above the graph represent p values.

To further investigate the consequences of infection, we performed H&E staining on lung tissue collected at day 7 post-infection. This time point was selected because the virus has recently been cleared, allowing us to assess residual tissue damage, cellular infiltrates, and structural changes. Histological analysis revealed distinct patterns across the three variants (**Figure 4b**). MA10-infected lungs exhibited focal inflammation, primarily near airways and vascular structures. Beta-infected lungs showed widespread tissue damage, including extensive alveolar and bronchial involvement and significant tissue consolidation. Omicron-infected lungs exhibited mild inflammation with a diffuse pattern, less extensive than Beta and more evenly distributed than the focal inflammation observed in MA10. Despite the limited number of infected cells, minimal PenH changes, and relatively mild histological damage observed in Omicron-infected lungs, some residual tissue consolidation was still detected at day 7. Consolidation scores, determined by unbiased quantification of open airspace versus stained tissue, revealed that Omicron- and MA10-infected lungs had similar scores at this timepoint (**Figure 4c**). In contrast, Beta infection significantly increased consolidation scores, consistent with severe lung damage and inflammation (**Figure 4c**). These findings highlight the distinct pathological profiles of the three variants and suggest that Omicron induces a milder lung pathology compared to Beta.

To further complement these findings, we analyzed the consequences of serial passaging in WT and *Ifitm3^-/-^*mice, focusing on the progression of adaptation between passage 10 and passage 20 Beta and Omicron viruses. These analyses revealed that Beta passage 20 viruses exhibited higher nasal and heart titers, greater airway dysfunction as measured by plethysmography, and more severe lung pathology compared to passage 10 viruses (**Supplementary Figure 5**). Omicron viruses showed consistent nasal titers for KO passages 10 and 20, consistently detectable heart titers by passage 20, minimal airway resistance measurements, and milder lung inflammation and structural damage relative to Beta (**Supplementary Figure 6**). Together, these results highlight how serial passaging in different host environments drives progressive adaptation of Beta and Omicron viruses, with Beta acquiring traits that exacerbate pathogenicity and Omicron maintaining features associated with reduced disease severity.

We compared lung inflammatory cytokine levels to gain insight into variant-specific immune responses (**Figure 4d**). Beta-infected mice exhibited the highest average levels of pro-inflammatory cytokines IFNβ, IL-1β, TNF, and IL-6. Omicron elicited moderate cytokine responses, except for IFNb, which was similarly strong to that induced by Beta. MA10 induced the lowest levels of all the inflammatory cytokines. These results suggest that the different variants may trigger the immune system through distinct mechanisms or that they may have differing mechanisms of immune evasion. In all, lung inflammatory cytokine levels were not an ideal predictor of lung dysfunction and pathology, suggesting additional mechanisms are active in driving variant-specific lung pathology.

Given the adaptation of Beta and Omicron strains in C57BL/6 mice, we next tested their pathogenicity in Balb/c mice to assess the influence of host genetic background. Distinct from its effects in C57BL/6 mice, Omicron KO passage 20 caused no detectable weight loss or changes in PenH (**Supplementary Figure 7a,b**), indicating minimal pathogenicity in the Balb/c genetic background. Differences in weight loss seen for Beta and MA10 strains in C57BL/6 mice were lost in Balb/c mice (**Supplementary Figure 7a**) though Beta continued to cause higher average PenH values throughout the course of infection in the Balb/c animals (**Supplementary Figure 7b**). These findings are consistent with the origins of MA10 as a Balb/c-adapted strain, while our Beta and Omicron variants were adapted in C57BL/6 mice. Together, these data underscore that host genetic background is critical in determining variant pathogenicity and disease outcomes.

### Defining the host transcriptional responses to adapted SARS-CoV-2 variants in the infected lung

To gain deeper insights into the mechanisms driving variant-specific pathology, we profiled the lung transcriptional responses to viral infection with MA10 or Passage 20 of Beta and Omicron from both WT and *Ifitm3^-/-^*mice at day 2 post-infection. This time point represents peak viral replication and has been associated with robust transcriptional responses to SARS-CoV-2 infection in mice^32,46,47,57^. Multidimensional scaling of the host transcriptional signatures revealed distinct responses elicited across viral strains with a lesser contribution of WT or KO passaging on the host response to infection (**Figure 5a**). Overall, MA10 and both Beta strains induced a greater magnitude of change in gene expression (**Figure 5b and Supplementary Figure 8a**), with a higher number of significant differentially expressed genes across these strains, relative to Omicron-induced gene expression changes (**Figure 5c**). Analysis of the overlap in host responses to infection identified 63 genes upregulated across all infectious conditions (**Figure 5d**) corresponding to interferon responses (**Supplementary Figure 8b**). Gene Ontology analysis confirmed the conservation of this crucial antiviral response^58–60^ across strains (**Figure 5e**).

**Figure 5.**
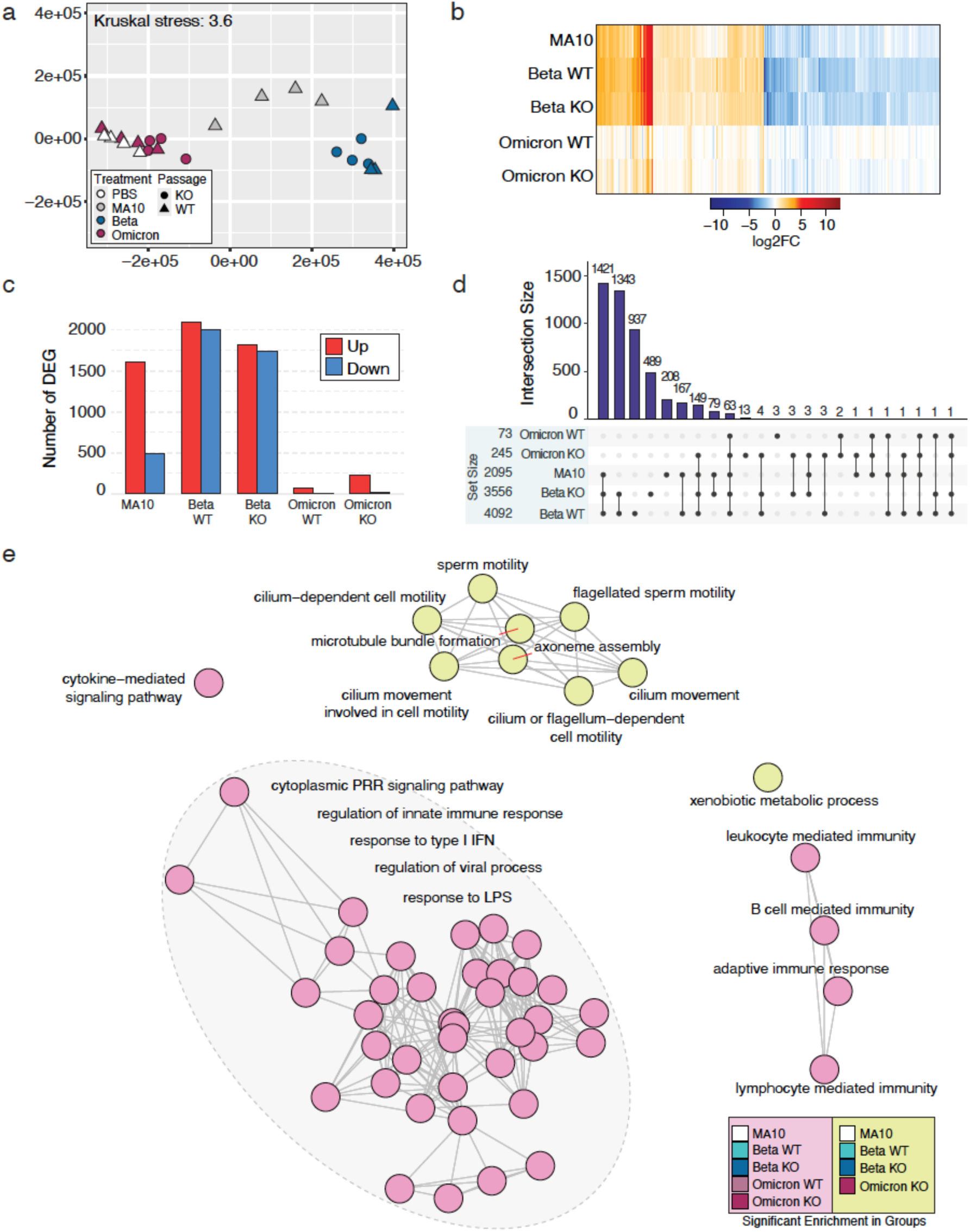
SARS-CoV-2 variants elicit distinct lung gene expression profiles at day 2 post-infection. (**a**) Multidimensional scaling (MDS) representation of the lung transcriptional responses elicited by SARS-CoV-2 mouse-adapted strains. Each data point represents a biological replicate, where color denotes viral and shape passage in WT or IFITM3 KO mice. The quality of the representation is provided by the Kruskal stress value, with a low percentage of Kruskal stress (3.6%). (**b**) Euclidean distance-based hierarchical clustering of the union (4,894 genes) of differentially expressed genes (DEG; lfc = |1|, padj <0.05). The heat map represents the average expression intensities for each infectious condition relative to the average of mock-infected samples. (**c**) Bar graphs represent the average numbers of genes upregulated (red) or downregulated (blue) in response to infection with the indicated viral strains. (**d**) Upset plot indicates the overlap of DEG across infectious conditions. (**e**) The affiliation network represents gene connectivity across GO Biological Pathways terms identified by Gene Set Enrichment Analysis (GSEA). Each node represents a unique term, edges represent connectivity across terms. Color indicates significant enrichment across infectious groups as indicated by color scale.

To further dissect these responses, we conducted a Weighted Gene Co-Expression Network Analysis with the union of differentially expressed genes across all conditions (**Supplementary Figure 9a**). This identified five key gene expression modules (Blue, Brown, Green, Turquoise, Yellow), with eigengenes summarizing their activities (**Table 1**). Unassigned genes were grouped under a ‘Grey’ module. Eigengenes are representative genes that summarize the expression profiles of gene modules identified through co-expression network analysis. We examined module eigengene expression to identify key patterns of expression across infectious conditions (**Supplementary Figure 9b**) and the relationship between modules (**Supplementary Figure 9c**). This revealed eigengenes predominantly upregulated (blue) or downregulated (Brown and Turquoise) across MA10 and Beta infections. Eigengenes within the yellow module appeared to be predominantly active in Beta infected lungs (**Supplementary Figure 9b**). To better understand the relationship between the features of viral disease and module eigengene expression, we calculated the area under the curve (AUC) for weight data across viral infections to identify significant differences between specific viral groups (**Figure 6a, left**). Correlation tests between module eigengenes and mean AUC identified a significant relationship between the eigengenes in the Brown module and disease severity (**Figure 6a, right**). Similarly, we calculated the AUC for lung function data (PenH) across viral infections and identified a significant increase in the PenH AUC in Beta KO and a significant decrease in PenH AUC in Omicron WT and KO-infected mice relative to MA10 (**Figure 6b, left**). Correlation tests between module eigengenes and mean AUC identified significant relationships between the eigengenes in the Brown and Turquoise module and disease severity (**Figure 6b, right**).

**Figure 6.**
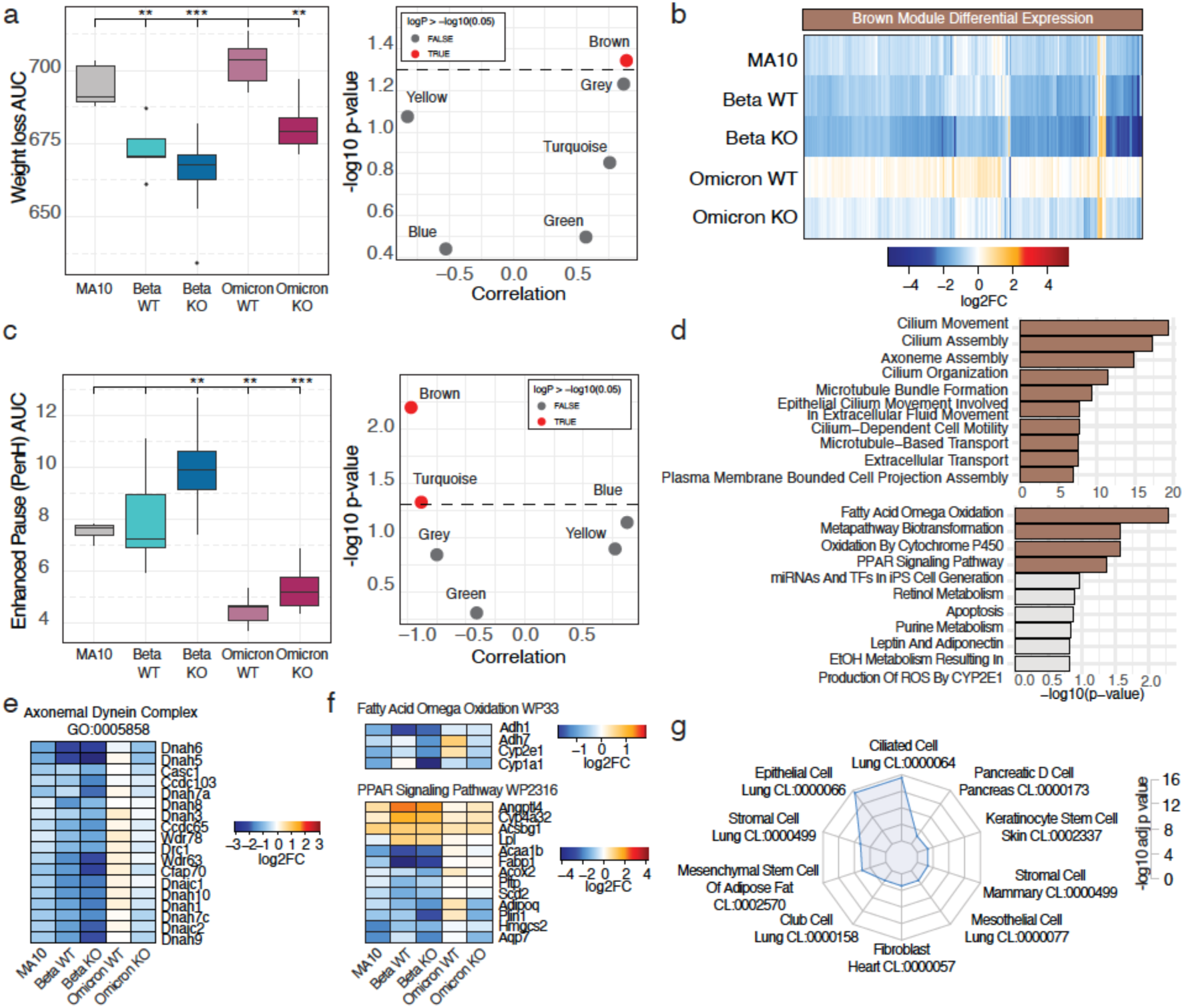
Network analysis reveals gene expression modules associated with viral disease. (**a**) Area under the curve (AUC) analysis of weight loss induced overtime by infection with SARS-CoV-2 mouse-adapted strains (left). Volcano plots representing the Pearson’s correlation coefficient (x axis) and -log10 significance value of gene expression modules with weight loss AUC. Dots represent WGCNA-identified modules and color indicates significant (red) or non-significant (grey) correlations. (**b**) Area under the curve (AUC) analysis of PenH induced over time by infection with SARS-CoV-2 mouse-adapted strains (left). Volcano plots representing the Pearson correlation (x axis) and -log10 significance value of gene expression modules with PenH AUC. Dots represent WGCNA-identified modules and color indicates significant (red) or non-significant (grey) correlations. (**c**) Euclidean distance-based hierarchical clustering of the 324 genes contained within the ‘Brown’ module. The heat map represents the average expression intensities for each infectious condition relative to the average of mock-infected samples. Color indicates genes upregulated (red) or downregulated (blue) in response to infection relative to mock-infected animals. (**d**) Bargraphs represent the -log10 padj significance values of the top 10 GO Biological Pathways (top) and WikiPathways Mouse (bottom) terms identified in the ‘brown’ module using EnrichR. (**e**) Heatmap representation of the average expression intensities across infectious condition relative to the average of mock-infected samples of DEG within the Axonemal Dynein Complex GO Biological Pathways term. Color indicates genes upregulated (red) or downregulated (blue) in response to infection relative to mock-infected animals. (**f**) Heatmap representation of the average expression intensities across infectious condition relative to the average of mock-infected samples of DEG within the Fatty Acid Omega oxidation (top) and PPAR Signaling Pathway (bottom) WikiPathways terms. Color indicates genes upregulated (red) or downregulated (blue) in response to infection relative to mock-infected animals. (**g**) Radar plots indicating the -log10 padj values of enrichment for Tabula Muris cell ontology terms within the 1343 DEG induced by mouse-adapted SARS-CoV-2 Beta strains.

We visualized changes in the expression of genes within the Brown module (**Figure 6c**) and noted a predominant decrease in the expression of genes involved in the function of cilia (**Figure 6d, top**). Genes involved in the axoneme dynein complex, the molecular motor required for the beating of cilia, were found to be predominantly downregulated in the lungs of mice infected with Beta passage 20 from WT or KO mice (**Figure 6e**). We also observed downregulation of genes related to fatty acid oxidation (**Figure 6d, bottom** and **Figure 6f, top**), a metabolic pathway known to be perturbed in alveolar epithelial cells in acute lung injury^61^. Viral infection also elicited pronounced changes in genes involved in PPAR signaling pathways (**Figure 6d, bottom** and **Figure 6f, bottom**) that are necessary to maintain barrier integrity and control lung inflammation^62,63^. Cell deconvolution enrichment analysis of the 1343 genes DEG unique to Beta WT and KO passage 20 infections (**Figure 6d**) revealed a predominant enrichment for ciliated cells, stromal cells, and club cells of the lung amongst others (**Figure 6g**). These data suggest increased barrier disruption elicited by Beta infection relative to MA10 and Omicron and are supported by histological examinations at days 2 and 7 post infection (**Figures 3e, 4b,c, and Supplementary Figures 5d, 6d**).

To better understand the biological functions of genes contained within the Turquoise module that correlated with PenH lung dysfunction measurements, we first examined their gene expression patterns and found genes that were increased and decreased in expression (**Figure 7a**). These genes were predominantly associated with the remodeling of the extracellular matrix and ion transport (**Figure 7b,c**). More specifically, the genes upregulated in the infected lungs (Orange, Yellow, and Back heatmap clusters) corresponded to genes involved in cytokine regulation, leukocyte migration, iron homeostasis, and non-canonical NF-kB signaling pathways (**Supplementary Figure 10a, left and 10c**). The genes downregulated in the infected lungs (Skyblue and Red heatmap clusters) corresponded to genes enriched in extracellular membrane organization (**Supplementary Figure 10a, right and 10d**). These findings were also consistent with a pronounced increase in expression of matrix metalloproteinases such as *Mmp10* and its inhibitor *Timp1* in MA10 and Beta, but not Omicron, infected lungs (**Figure 7d**). The expression of *Mmp8* (neutrophil collagenase) and *Mmp25* (leukolysin), which are produced primarily by neutrophils^64–66^, was also increased predominantly in Beta infections and to some extent in MA10 infected lungs.

**Figure 7.**
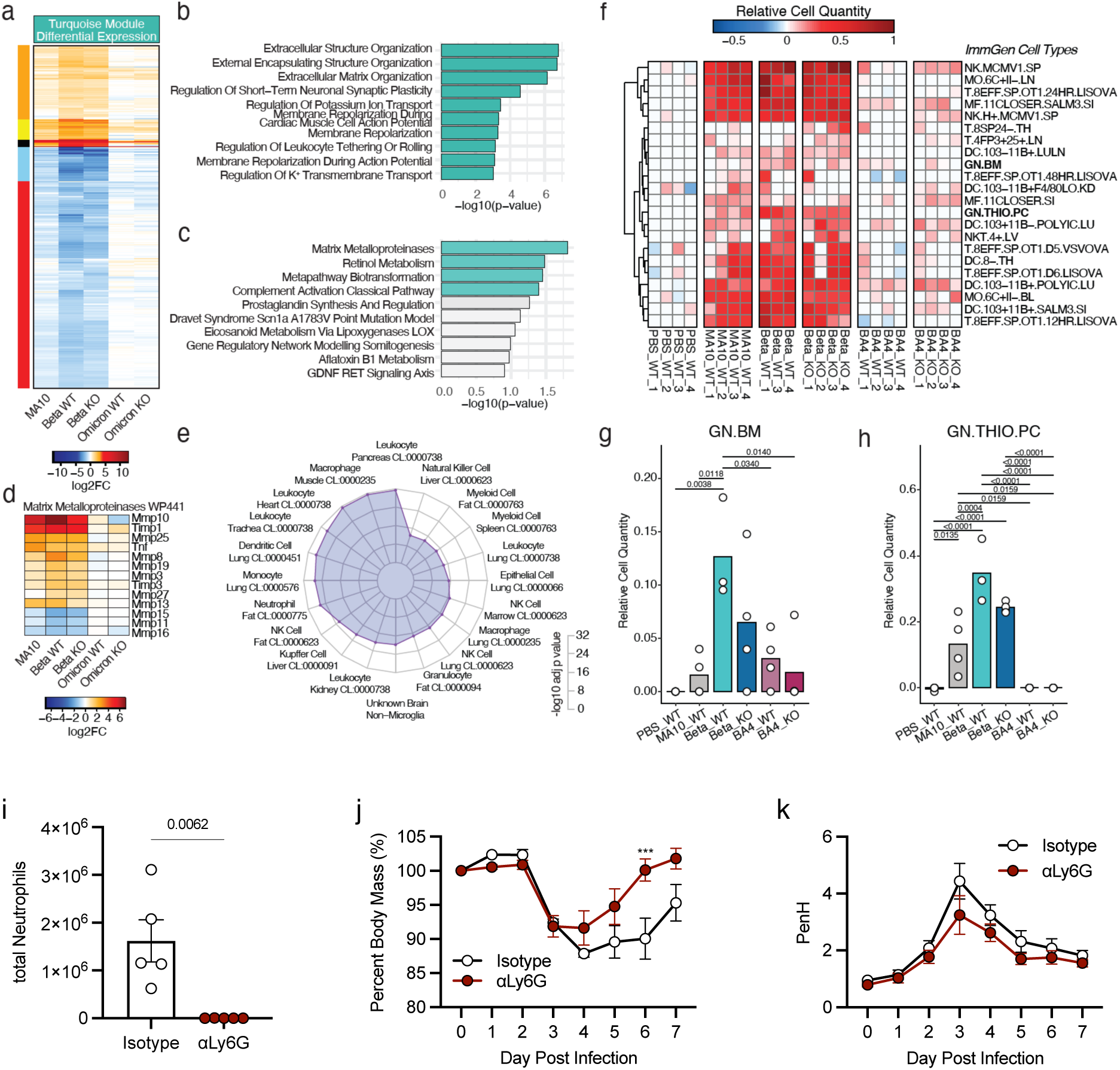
Excessive monocyte infiltration correlates with disruption in lung function. (**a**) Euclidean distance-based hierarchical clustering of the 2342 genes contained within the ‘Turquoise’ module. The heat map represents the average expression intensities for each infectious condition relative to the average of mock-infected samples. Color indicates genes upregulated (red) or downregulated (blue) in response to infection relative to mock-infected animals. (**b**) Bargraphs represent the -log10 padj significance values of the top 10 GO Biological Pathways terms identified in the ‘Turquoise’ module using EnrichR. (**c**) Bar graphs represent the -log10 padj significance values of the top 10 WikiPathways Mouse terms identified in the ‘Turquoise’ module using EnrichR. (**d**) Heatmap representation of the average expression intensities across infectious condition relative to the average of mock-infected samples of DEG within the Matrix Metalloproteinases WikiPathway term. Color indicates genes upregulated (red) or downregulated (blue) in response to infection relative to mock-infected animals. (**e**) Radar plots indicating the -log10 padj values of enrichment for Tabula Muris cell ontology terms within the 1343 DEG induced by mouse-adapted SARS-CoV-2 Beta strains. (**f**) DCQ inference of immune cell subsets derived from whole-lung gene expression across infectious conditions. Each row represents an ImmGen cell type. Columns indicate individual infectious conditions. Color indicates positive (red) and negative (blue) relative cell quantity values. Only positively enriched cell types are shown. (**g-h**) Bar graphs representing DCQ relative cell quantity values for the ImmGen granlulocyte populations (**g**) GN.BM and (**h**) GN.THIO.PC. Significance determined by one-way ANOVA with Tukey HSD pairwise comparison test. (**i-k**) Groups of WT mice (n=5 per group) were challenged with 10^5^ TCID50 of the SARS-CoV-2 Beta strain passaged 20 times through *Ifitm3^-/-^* mice. To deplete neutrophils, mice were treated with antibodies targeting neutrophils (αLy6G) or an isotype control antibody, starting at day 1 post infection through day 6 post infection. (**i**) Neutrophil number in the lungs of infected mice on day 4 post infection by flow cytometry; error bars indicate SEM, p value above the bar determined by two-tailed unpaired t-test. (**j**) Weight loss during challenge by passaged SARS-CoV-2 viruses. Error bars represent SEM, comparisons were made using the Mann-Whitney test. (**k**) Daily whole-body plethysmography enhanced pause (PenH) measurements. (**j, k**) Dots represent averages of individual mice (n=5 per group).

Given the importance of these metalloproteases in the regulation of angiogenesis and inflammation, we conducted cell deconvolution enrichment analysis to determine whether the shared gene signatures induced by MA10 and Beta SARS-CoV-2 infections (1421 DEG, **Figure 1d**) could predict changes in distinct cell subsets within infected lungs. These gene signatures revealed an enrichment of leukocyte, macrophage, dendritic cell, and neutrophil cell subsets (**Figure 7e**). We corroborated this inference using digital cell quantifier (DCQ), a quantitative deconvolution tool that has been validated using flow cytometric measurements^67^ and that we have previously used to predict immune cell infiltration following respiratory pathogen infection^68,69^. DCQ inferred high positive enrichment of activated immune cell types, including NK cells, CD8+ T cells, macrophages, and dendritic cells, in the lungs of mice infected with MA10 and Beta, but not Omicron (**Figure 7f**). Negatively enriched gene signatures corresponded largely to hematopoietic and thymic precursors, reflecting the mature, activated status of the immune cell types found in the lung following infection with MA10 or Beta (**Supplementary Figure 10e**). Interestingly, we noted that two granulocyte populations, GN.BM and GN.THIO.PC, were most significantly enriched in Beta infection relative to mock-inoculated, MA10, and Omicron infections (**Figure 7f-h and Supplementary Figure 10 f-g**). These data suggest that neutrophil infiltration could contribute to disease severity in SARS-CoV-2 infections.

Given that neutrophilia is highly associated with increased disease severity in human SARS-CoV-2 infections^70–75^, and given that we and others previously showed robust recruitment of neutrophils to MA10-infected mouse lungs^46,57,76^, we examined the impact of antibody-mediated neutrophil depletion on disease caused by Beta KO passage 20, our most virulent viral strain. Confirming the inferences from our transcriptomic analysis, neutrophil depletion significantly decreased weight loss and time to recovery and accelerated recovery to baseline PenH readings (**Figure 7i-k**). Together, these findings reveal how variant-specific host transcriptional responses, particularly the regulation of cilia function, extracellular matrix remodeling, and neutrophil recruitment, underpin the pathological outcomes of SARS-CoV-2 infections.

## DISCUSSION

Our study provides critical insights into the role of IFITM3 in restricting the interspecies adaptation of SARS-CoV-2 and highlights distinct pathogenic characteristics of Beta and Omicron variants. We demonstrated that the absence of IFITM3 facilitates the accumulation of adaptive mutations upon passaging of human-derived virus in mice, allowing it to more rapidly overcome barriers to replication and dissemination in the new host species. These findings underscore the potential for IFITM3 to act as a key antiviral effector during zoonotic transmission and emphasize how deficiencies in this protein could influence the evolution of emerging pathogens, particularly given the prevalence of IFITM3 defects in humans^23,26–31,77,78^.

The generation of new mouse-adapted Beta and Omicron variants uniquely enabled direct comparisons of their distinct replication and pathogenic profiles in a controlled model. This approach allowed us to dissect which effects are intrinsic to the genetic foundation of each strain, independent of their original adaptation to human hosts. Beta exhibited severe lung pathology characterized by extensive spread of the virus from the large airways throughout the alveolar space. Beta infection was further associated with disruptions in cilia gene expression and extracellular matrix remodeling along with heightened inflammation and neutrophil gene signatures, each of which is consistent with its aggressive spread and virulence in humans. Indeed, these disruptions likely contribute to increases in airway resistance and weight loss observed in our studies. In contrast, Omicron infections displayed attenuated lung pathology, minimal airway dysfunction, reduced tissue inflammation, and heightened nasal titers, reflecting a preference for the upper respiratory tract in humans that was maintained upon mouse adaptation.

The use of transcriptional profiling provided additional mechanistic insights into the distinct host responses elicited by these variants. Downregulation of cilia-related genes and fatty acid oxidation pathways was particularly pronounced in Beta infections, consistent with histological evidence of airway damage and tissue consolidation. Pathways associated with PPAR signaling, which are critical for maintaining barrier integrity and controlling inflammation, were also significantly perturbed in Beta-infected lungs. These findings suggest that Beta induces severe lung pathology through a combination of direct viral damage and dysregulation of host metabolic and structural processes. By contrast, Omicron exhibited far fewer transcriptional changes in these pathways, which may contribute to its milder pathology.

The newly adapted Beta and Omicron variants also exhibited distinct patterns of extrapulmonary dissemination, with both strains detected in cardiac tissue, unlike MA10. This observation aligns with the known clinical association of SARS-CoV-2 variants with cardiac complications and underscores the utility of these mouse-adapted strains for studying the systemic effects of infection. Importantly, the adaptation of Beta and Omicron in C57BL/6 mice expands the range of available experimental models for SARS-CoV-2 research, enabling studies in genetically diverse or immunologically modified backgrounds. This represents a more severe model of infection in C57BL/6 mice compared to MA10, which was generated in Balb/c mice and has relatively mild pathogenicity in other mouse strains.

Interestingly, both Beta and Omicron variants carry the N501Y mutation in the Spike protein, which is known to allow binding to murine ACE2 and enable rodent infectivity^48^. This has led to the hypothesis that such variants may have evolved in rodents before being transmitted back to humans^79,80^. However, our findings challenge this scenario given that both Beta and Omicron variants exhibited poor initial replication and pathogenicity in mice, requiring serial passaging in the absence of IFITM3 to achieve robust adaptation. This suggests that while N501Y may confer a baseline ability to infect mice, the broader evolutionary context of these variants is unlikely to have involved sustained rodent hosts. Instead, the N501Y mutation has been shown to increase replication in primary human epithelial cells as well as increase the affinity of Spike to human ACE2^81^, suggesting that increased binding to murine ACE2 is an evolutionary byproduct. Our findings also have broader implications for understanding zoonotic transmission and pandemic risk. The accelerated adaptation observed in IFITM3-deficient mice highlights how host factors can influence the evolutionary trajectory of emerging viruses. Our data also indicate that IFITM3 deficiency does not necessarily promote unique mutations specific to its absence but rather accelerates the adaptive process that would have occurred under natural selective pressures. This highlights IFITM3 as a key host factor in slowing the rate of viral adaptation without fundamentally altering the trajectory of mutation acquisition. This suggests that populations with genetic or acquired deficiencies in IFITM3 may serve as permissive environments for zoonotic coronaviruses to adapt to human hosts. This knowledge should inform interventions for IFITM3-deficient individuals to reduce the risk of zoonotic spillover and the emergence of adapted viral strains.

In conclusion, this study establishes the critical role of IFITM3 in limiting the adaptation and pathogenesis of SARS-CoV-2 variants and provides new tools to dissect the molecular and immunological determinants of variant-specific disease. The distinct biological properties of the adapted Beta and Omicron strains emphasize the complex interplay between host and viral factors that drives pathogenicity, overall advancing our understanding of SARS-CoV-2 biology.

## RESOURCE AVAILABILITY

All data are available in the main text or the supplementary materials. Requests for further information and resources should be directed to and will be fulfilled by the lead contact, Jacob Yount.

## ACKNOWLEDGMENTS

The authors thank the Genomics and Microbiology Solutions Laboratory (IDI-GEMS, OSU) for providing virus sequencing and analysis support. We would like to thank Allison Sasso and JA Fortou for helpful discussion. Research in the Yount Laboratory is funded by NIH grants AI130110, HL168501, HL157215, HL154001, and AI151230. JR is supported by an American Heart Association Graduate Research Fellowship. Research in the Forero Laboratory is funded by NIH grants GM150806 and T32AI165391 (MIM).

## AUTHOR CONTRIBUTIONS

P.J.D. organized and performed all experiments in the manuscript, analyzed data, and contributed to manuscript writing, figure generation, and editing of the manuscript. J.L.P. and J.R. assisted with mouse colony genotyping/maintenance, virus titering, and mouse experiments. M.I.M. and P.R. R. assisted with analysis of RNA sequencing data and contributed to manuscript writing and figure generation. A.F. provided conceptual input, experimental design, RNA sequencing analysis, interpreted data, and wrote the manuscript. J.S.Y. conceived the study, supervised experimental design and performance of experiments, analyzed and interpreted data, and wrote the manuscript.

## DECLARATION OF GENERATIVE AI AND AI-ASSITED TECHNOLOGIES

During the preparation of this work the authors used ChatGPT in order to improve language and readability of this manuscript. After using this tool, the authors reviewed, edited, and incorporated suggested changes as needed and take full responsibility for the content of the published article.

## SUPPLEMENTAL INFORMATION

Document S1. Figures S1–S10

## MATERIALS AND METHODS

### Biosafety

All experiments with SARS-CoV-2 were performed in the Ohio State University (OSU) Biosafety Level 3 (BSL3) facility. Experimental procedures were approved by the OSU Institutional Biosafety Committee and OSU BSL3 Advisory Group under protocol number 2020R00000025. Research samples analyzed outside BSL3 containment were decontaminated according to experimentally validated and institutionally approved protocols. Additional oversight for SARS-CoV-2 mouse adaptation experiments was provided by an *ad hoc* dual use research of concern evaluation committee assembled by the OSU Office of Research at the request of the principal investigator.

### Virus stocks and in vitro infection

All SARS-CoV-2 strains were obtained from BEI Resources including the Beta variant (hCoV-19/USA/MD-HP01542/2021, NR-55282), the BA.4 variant (hCoV-19/USA/MD-HP30386/2022; NR-56803), and mouse-adapted SARS-CoV-2 strain MA10 (generated by the laboratory of Dr. Ralph Baric, University of North Carolina; NR-55329). MA10 was plaque purified and sequenced prior to use. All viruses were propagated in Vero-TMPRSS2 cells (BEI Resources, NR-54970). Virus-containing supernatants were aliquoted, flash frozen in liquid nitrogen, and stored at −80°C. Viral stocks and research samples were titered via TCID50 measurements on Vero E6 (ATCC, CRL-1586) cells using 10-fold serial dilutions in triplicate. The presence of virus replication in each well was assessed by visual examination of cytopathic effect along with antibody staining for N protein antigen (mouse monoclonal antibody 40143-MM08, Sino Biological) with fluorescent secondary antibody detection (Goat anti-Mouse IgG (H+L) Highly Cross-Adsorbed Secondary Antibody, Alexa Fluor™ 647 (Thermo Scientific, Catalog # A-21236)). TCID50 titers were calculated using the Reed–Muench method. Cells were maintained in DMEM medium (Fisher scientific) supplemented with 10% EquaFETAL bovine serum (Atlas Biologicals). All cells were cultured in a humidified incubator at 37 °C with 5% CO2. Cell lines were confirmed mycoplasma-negative prior to use.

### Mouse studies

*Ifitm3^-/-^* mice on the C57BL/6 genetic background with a 53 base pair deletion in exon 1 of the *Ifitm3* gene were described previously^33^. Mice were housed in the OSU Biomedical Research Tower ABSL3 vivarium, which is maintained at 68–76 degrees F, with a 12:12 light dark cycle, and humidity between 30–70%. Autoclaved individually ventilated cages (Allentown) were used for housing. Mice were fed irradiated natural ingredient chow ad libitum (Evnigo Teklad Diet 7912). Reverse osmosis purified water was provided through an automated rack water system. Cages included 0.25 inch of corn cob bedding (Bed-o-Cobs, The Andersons) with cotton square nesting material. Male and female mice between 6 and 8 weeks of age were used in our experiments. All mice in each individual experiment were age-matched. All mouse infections were performed intranasally under anesthesia with isoflurane, delivering 25μL of inoculum per nostril. For virus passaging, mice were initially infected with doses of 10^5^ TCID50, and mouse organs were collected and homogenized in 500μL of sterile PBS. Propagation and titration of passaged viruses, as well as parental control viruses, was done on Vero-TMPRSS2 cells. WT C57BL/6 mice (Charles River Laboratories) were infected with equal doses (10^5^ TCID50) for testing of virus adaptation. Innoculums were prepared via dilution of viral stock in sterile saline. Mouse organs were collected and homogenized in PBS for ELISAs or titration, homogenized in Trizol for extraction of RNA, or fixed in 10% formalin for histology. All procedures were approved by the OSU IACUC (protocol 2020A00000054-R1) and were performed following guidelines for the ethical use of animals.

### ELISAs

ELISAs were performed on mouse lung homogenates collected at day 2 post-infection. All lungs were homogenized in 1 mL sterile PBS and centrifuged prior to supernatant collection. Assays were performed using the Mouse IL-6 Duoset ELISA kit from R&D Systems (catalog # DY406), the Mouse IFNβ Duoset ELISA kit from R&D Systems (catalog # DY8234-05), the Mouse IL-1β Duoset ELISA kit from R&D Systems (catalog # DY401), or the Mouse TNFα Duoset ELISA kit from R&D Systems (catalog # DY410) according to manufacturer’s instructions.

### Histology

For lung histology, lung tissue samples were fixed in 10% formalin for 7 days at 4°C and subsequently transferred to 70% ethanol. Lungs were sectioned, stained, and imaged by Histowiz (<u>Histowiz.com</u>, Brooklyn, NY, USA). Quantification of the H&E images was performed using ImageJ software and the color deconvolution method^32,33,57,82^. Lung sections were stained with an anti-SARS-CoV-2 N protein antibody (Histowiz, #GTX635686) IHC images were analyzed using ImageJ software to quantify the DAB staining of lung tissue as previously described^32,83^. For analyses of each sample, whole lung images were analyzed via the color deconvolution method.

### RNA Sequencing

RNA was extracted from lung tissue homogenates using TRIzol (Invitrogen). Following separation with 1-bromo-3-chloropropane, aqueous phase was collected, and RNA was precipitated with ethanol. Precipitated RNA was isolated and sent to GENEWIZ, LLC/Azenta US, Inc (South Plainfield, NJ, USA). RNA library preparation and sequencing was performed with the following methods as provided by GENEWIZ: “The RNA samples received were quantified using Qubit 2.0 Fluorometer (ThermoFisher Scientific, Waltham, MA, USA) and RNA integrity was checked using TapeStation (Agilent Technologies, Palo Alto, CA, USA). The RNA sequencing libraries were prepared using the NEBNext Ultra II RNA Library Prep Kit for Illumina using manufacturer’s instructions (New England Biolabs, Ipswich, MA, USA). Briefly, mRNAs were initially enriched with Oligod(T) beads. Enriched mRNAs were fragmented for 15 min at 94°C. First strand and second strand cDNA were subsequently synthesized. cDNA fragments were end repaired and adenylated at 3′ends, and universal adapters were ligated to cDNA fragments, followed by index addition and library enrichment by PCR with limited cycles. The sequencing libraries were validated on the Agilent TapeStation (Agilent Technologies, Palo Alto, CA, USA), and quantified by using Qubit 2.0 Fluorometer (ThermoFisher Scientific, Waltham, MA, USA) as well as by quantitative PCR (KAPA Biosystems, Wilmington, MA, USA). The sequencing libraries were multiplexed and clustered onto a flowcell. After clustering, the flowcell was loaded onto the Illumina HiSeq instrument according to manufacturer’s instructions. The samples were sequenced using a 2 × 150 bp Paired End (PE) configuration. Image analysis and base calling were conducted by the HiSeq Control Software (HCS). Raw sequence data (.bcl files) generated from Illumina HiSeq was converted into fastq files and de-multiplexed using Illumina bcl2fastq 2.17 software. One mis-match was allowed for index sequence identification.”

Sequencing alignment and gene counts were generated using Rosalind as previously described^32,84^. Quality control of processed count data for outlier assessment, filtering of low expressed genes, and normalization was done using R version 4.4.2. Statistical analysis for differential gene expression was done using DESeq2 (v 1.46.0). Network analysis was performed using ‘WGCNA’ (v 1.73) and functional analysis was done using ‘enrichR’ (v 3.2) in R. Functional enrichment and cell deconvolution was performed with ‘clusterProfiler’, ‘msigdbR’, ‘fgsea’, ‘enrichR’ packages for R. Digital cell quantifier (DCQ) analysis^85^ was performed from genome-wide lung transcriptional profiles using ‘ComICS’ as we have previously done^68,69^. Data visualization was performed in R.

Statistical Analysis

Area under the curve (AUC) analysis was calculated using the trapezoidal rule using the R package ‘pracma’ (v 2.4.4). Statistical significance (p <0.05) was calculated using ANOVA followed by Tukey’s Honestly Significant (HSD) post hoc analysis using R. All other statistical analyses were performed using GraphPad Prism software (v 10.4.1) with individual statistical tests indicated in the corresponding figure legends. P<0.05 was considered significant.

---

**Supplementary Figure 1.**
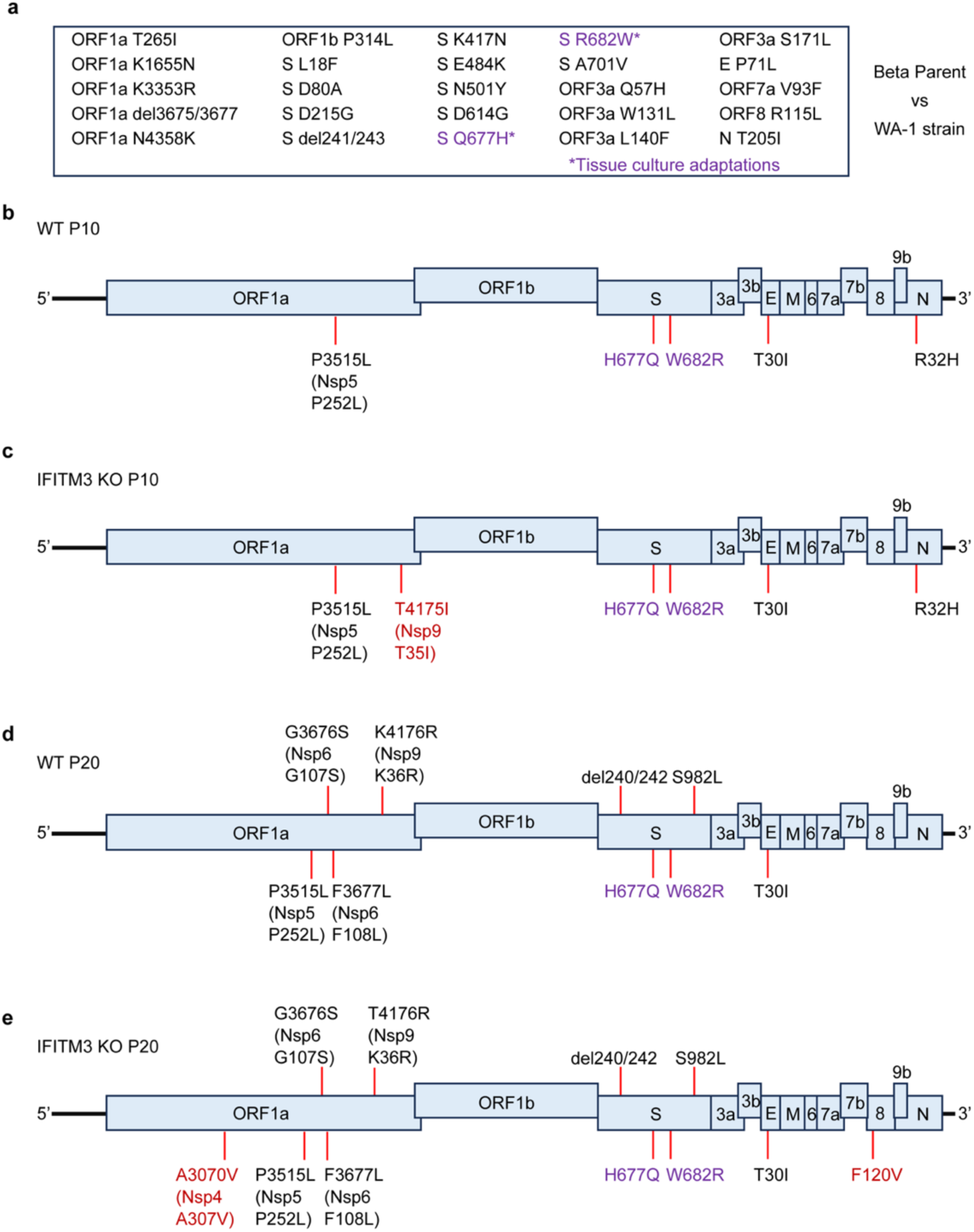
Mutations observed within the SARS-CoV-2 beta strain genome after passaging through WT or *Ifitm3^-/-^* mice. (**a**) List of amino acid differences between the SARS-CoV-2 Beta parent stock and the original WA1 strain. Tissue culture adaptations are indicated with an asterisk (*) and are colored purple. (**b, c**) Schematic of SARS-CoV-2 Beta strain genome annotated with amino acid substitutions found in the consensus sequence after passaging 10 times (**b**) or 20 times (**c**) through WT mice. (**d, e**) Schematic of SARS-CoV-2 Beta strain genome annotated with amino acid substitutions found in the consensus sequence after passaging 10 times (**d**) or 20 times (**e**) through *Ifitm3^-/-^* mice.

**Supplementary Figure 2.**
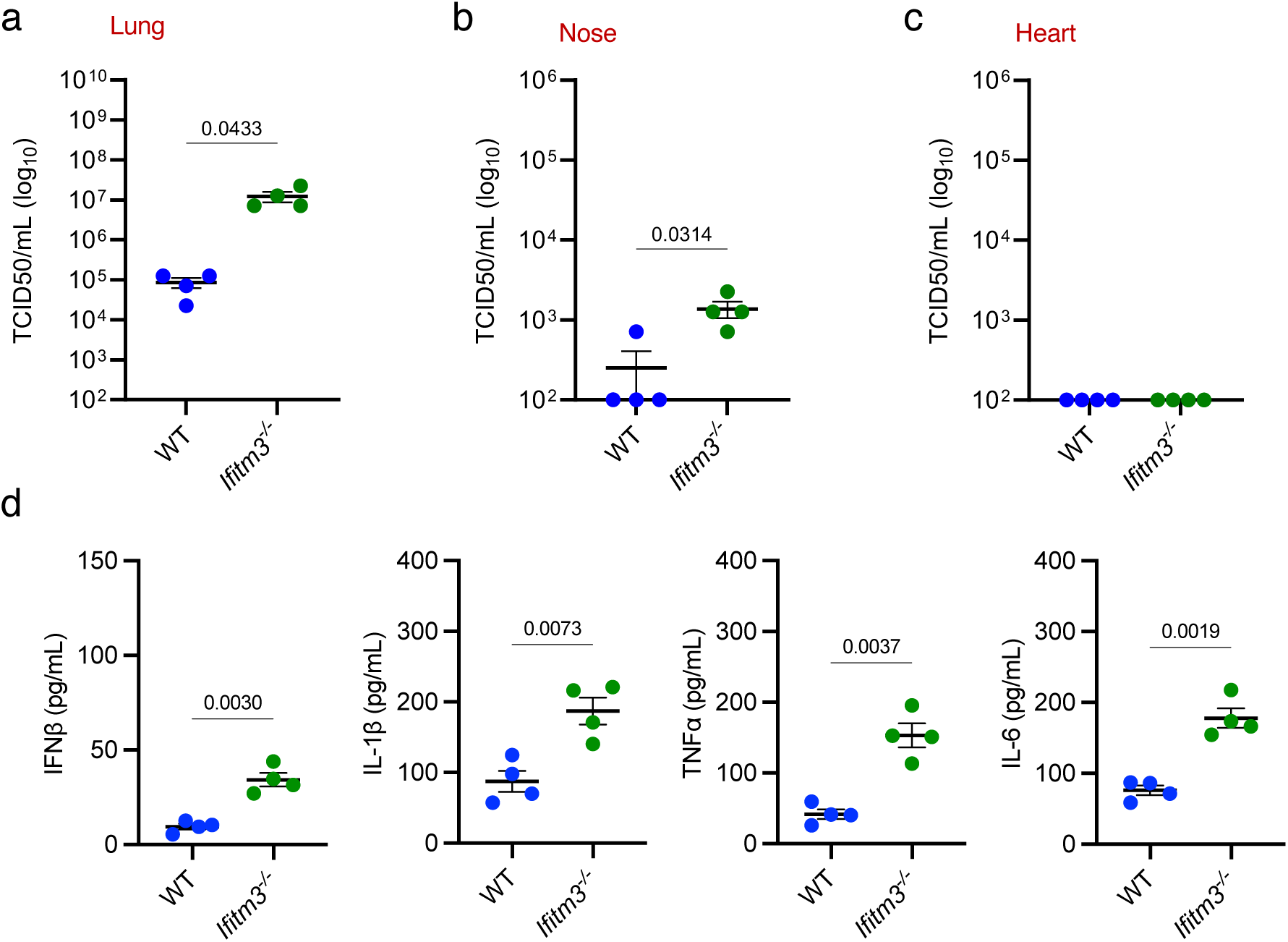
IFITM3 restricts the BA.4 omicron virus in mice. Groups of WT or *Ifitm3^-/-^*mice (n=4 per group) were challenged with 10^5^ TCID50 of the SARS-CoV-2 BA.4 strain. Viral titers from lung (**a**) or nose (**b**) homogenates collected at day 2 post infection. (**c**) ELISA quantification of IFNβ, IL-1β, TNFα, and IL-6 in lung homogenates collected at day 2 post infection. Each dot represents an individual mouse. Numbers above the graph represent p values.

**Supplementary Figure 3.**
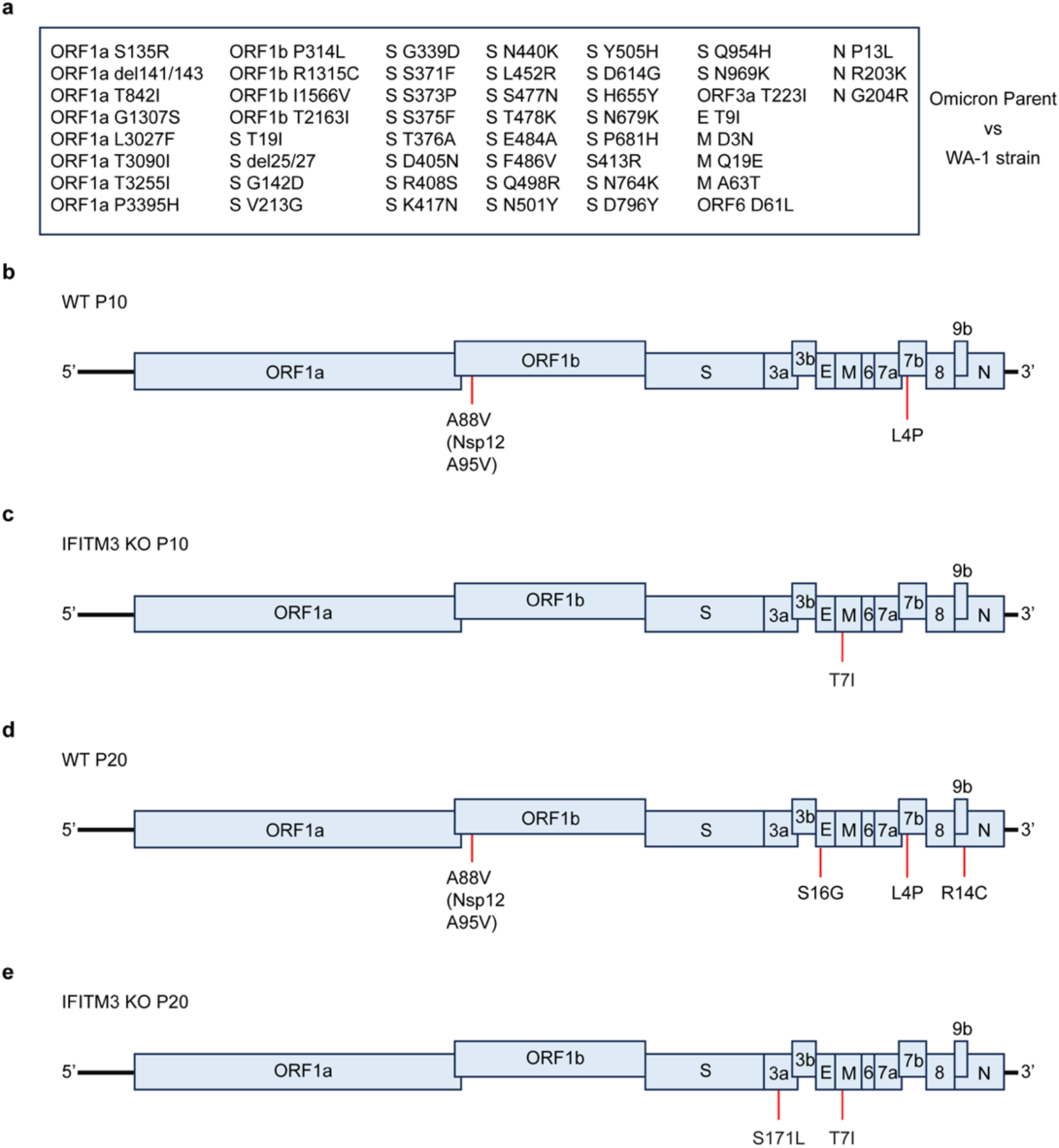
Mutations observed within the SARS-CoV-2 BA.4 strain genome after passaging through WT or *Ifitm3^-/-^* mice. **(a)** List of amino acid differences between the SARS-CoV-2 Beta parent stock and the original WA1 strain. (**b, c**) Schematic of SARS-CoV-2 BA.4 strain genome annotated with amino acid substitutions found in the consensus sequence after passaging 10 times (**b**) or 20 times (**c**) through WT mice. (**d, e**) Schematic of SARS-CoV-2 BA.4 strain genome annotated with amino acid substitutions found in the consensus sequence after passaging 10 times (**d**) or 20 times (**e**) through *Ifitm3^-/-^* mice.

**Supplementary Figure 4.**
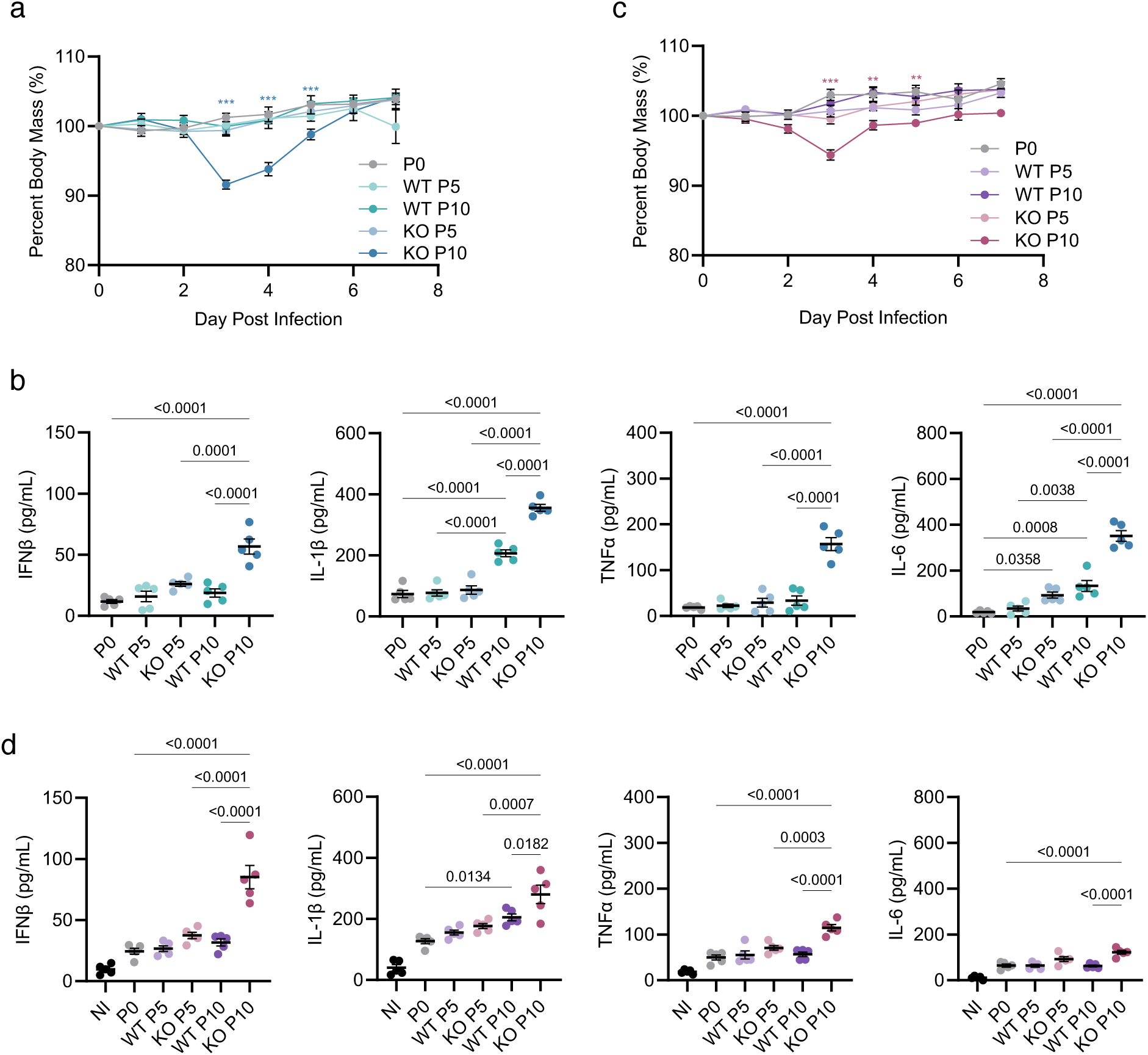
Passaging SARS-CoV-2 variants 10 times through *Ifitm3^-/-^* mice is sufficient to induce greater weight loss and inflammatory cytokines. Groups of WT mice (n=5 per group) were challenged with 10^5^ TCID50 of the SARS-CoV-2 Beta (**a, b**) or Omicron BA.4 (**c, d**) strains passaged 5 or 10 times through WT or *Ifitm3^-/-^* mice and compared to the parent virus (passage 0). (**a, c**) Weight loss during challenge by passaged SARS-CoV-2 viruses. Error bars represent SEM, comparisons were made using the Mann-Whitney test (only comparisons between P0, WT P10, and KO P10 are shown (p values: *** <0.0005, ** <0.005, *<0.05). Dots represent averages of individual mice (n=5 per group). (**b, d**) ELISA quantification of IFNβ, IL-1β, TNFα, and IL-6 in lung homogenates collected at day 2 post infection. Each dot represents an individual mouse. Numbers above the graph represent p values.

**Supplementary Figure 5.**
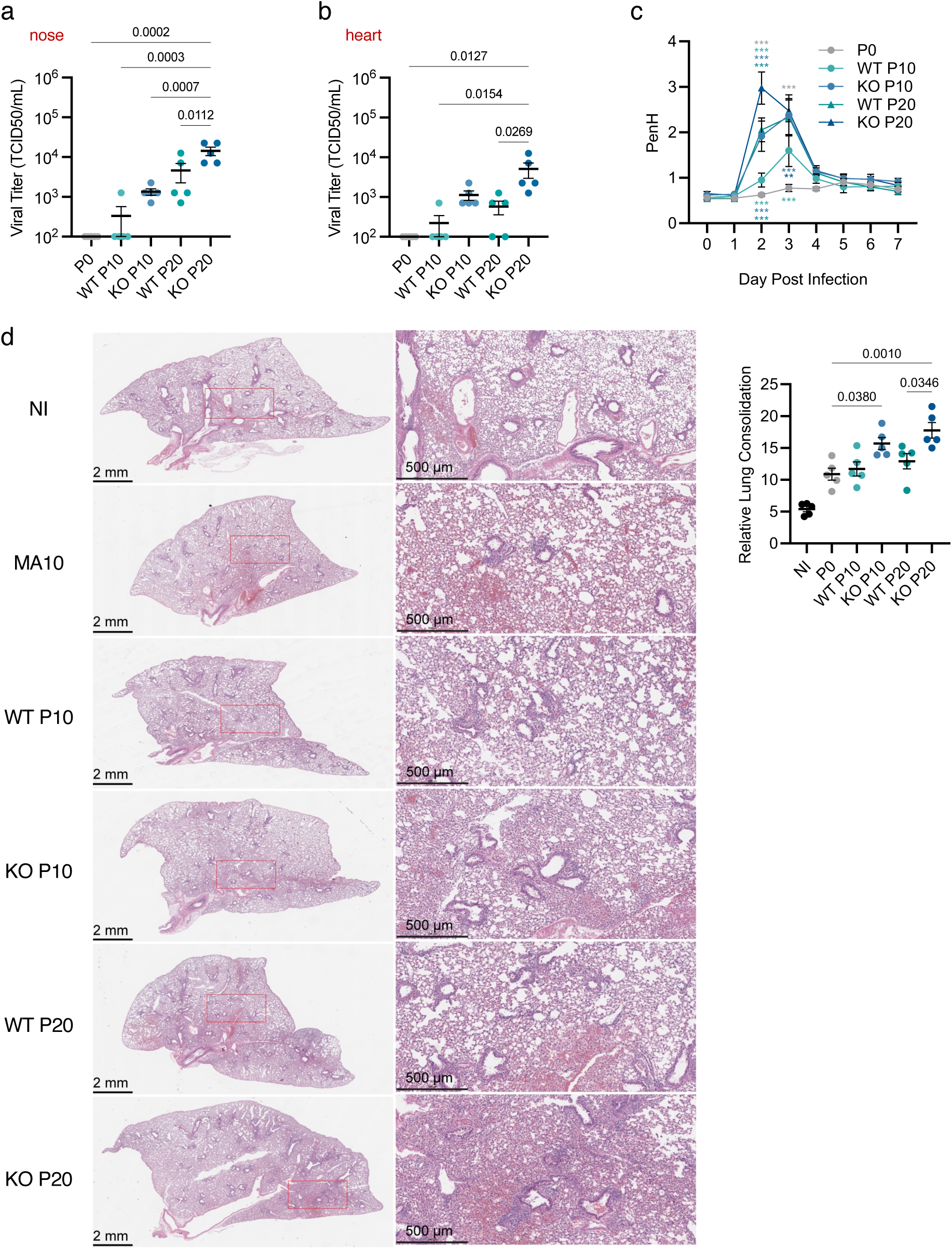
Comparing lung pathogenesis and inflammation induced by the SARS-CoV-2 Beta strain passaged 10 or 20 times through WT or *Ifitm3^-/-^* mice. Groups of WT mice (n=5 per group) were challenged with 10^5^ TCID50 of the SARS-CoV-2 Beta strain passaged 10 or 20 times through WT or *Ifitm3^-/-^* mice and compared to the parent virus (passage 0). Viral titers from nose (**a**), or heart (**b**) homogenates collected at day 2 post infection. (**c**) Daily whole-body plethysmography enhanced pause (PenH) measurements. Dots represent averages of individual mice (n=5 per group) (p values: **** <0.0001, *** <0.0005, ** <0.005, *<0.05). (**d**) Representative histology images of infected lungs at day 7 post infection. Whole lung images are shown on the left (scale bars represent 2 mm) and 4x zoomed images are shown on the right (scale bars represent 500 μm). The red box present in the whole lung images corresponds to its matched zoomed in image. (**e**) Quantification of consolidated tissue versus open airspace in H&E staining for entire lung sections of multiple animals infected at day 7 post infection. (**a, b, e**) Each dot represents an individual mouse. (**c**) Dots represent averages of individual mice (n=5 per group).

**Supplementary Figure 6.**
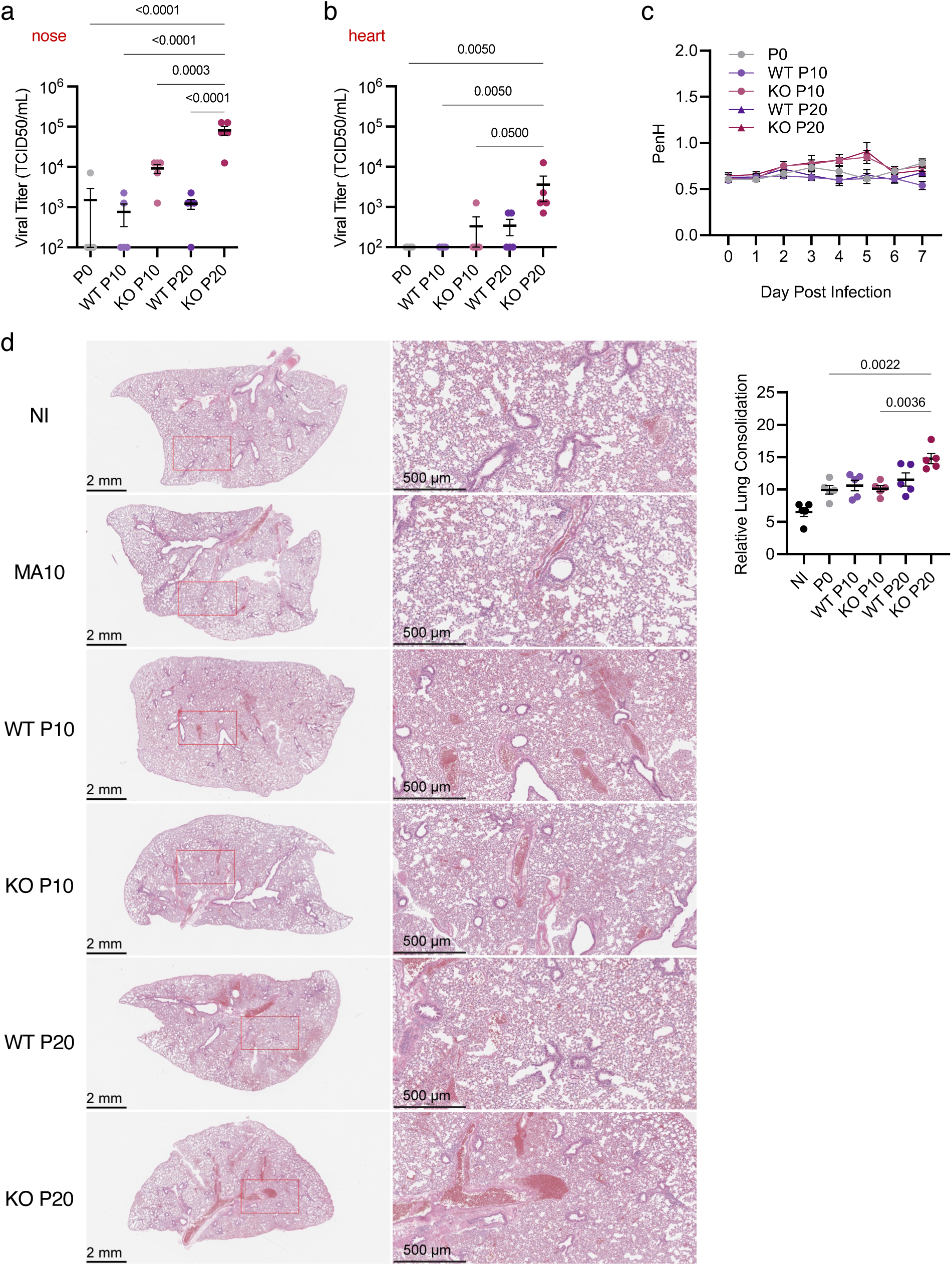
Comparing lung pathogenesis and inflammation induced by the SARS-CoV-2 Omicron BA.4 strain passaged 10 or 20 times through WT or *Ifitm3^-/-^* mice. Groups of WT mice (n=5 per group) were challenged with 10^5^ TCID50 of the SARS-CoV-2 BA.4 strain passaged 10 or 20 times through WT or *Ifitm3^-/-^*mice and compared to the parent virus (passage 0). Viral titers from nose (**a**), or heart (**b**) homogenates collected at day 2 post infection. (**c**) Daily whole-body plethysmography enhanced pause (PenH) measurements. Dots represent averages of individual mice (n=5 per group). (**d**) Representative histology images of infected lungs at day 7 post infection. Whole lung images are shown on the left (scale bars represent 2 mm) and 4x zoomed images are shown on the right (scale bars represent 500 μm). The red box present in the whole lung images corresponds to its matched zoomed in image. (**e**) Quantification of consolidated tissue versus open airspace in H&E staining for entire lung sections of multiple animals infected at day 7 post infection. (**a, b, e**) Each dot represents an individual mouse. (**c**) Dots represent averages of individual mice (n=5 per group).

**Supplementary Figure 7.**
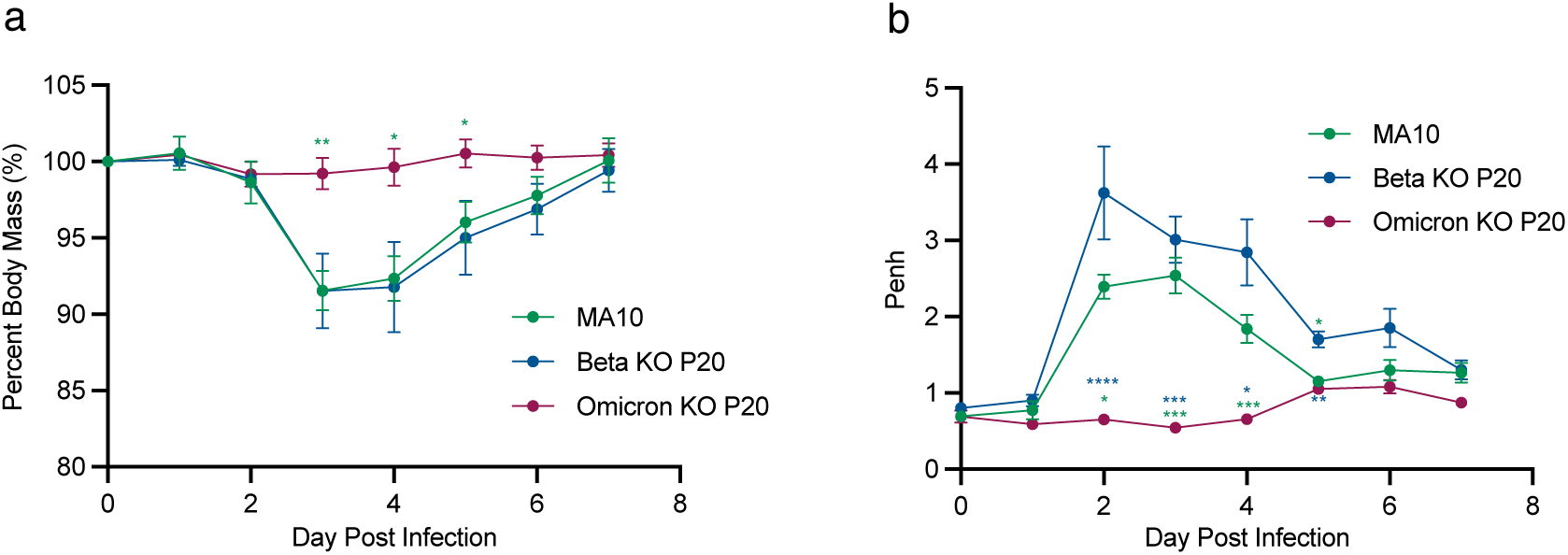
Comparative analysis of mouse-adapted Beta, Omicron, and MA10 strains in WT Balb/c mice. Groups of WT mice (n=5 per group) were challenged with 10^5^ TCID50 of the SARS-CoV-2 BA.4 strain or Beta strain passaged 20 times through *Ifitm3^-/-^* mice and compared to infection by an equal dose of MA10. (**a**) Weight loss during challenge by passaged SARS-CoV-2 viruses. (**b**) Daily whole-body plethysmography enhanced pause (PenH) measurements. (**a, b**) Dots represent averages of individual mice (n=5 per group). Error bars represent SEM, comparisons were made using the Mann-Whitney test. (p values: **** <0.0001, *** <0.0005, ** <0.005, *<0.05).

**Supplementary Figure 8.**
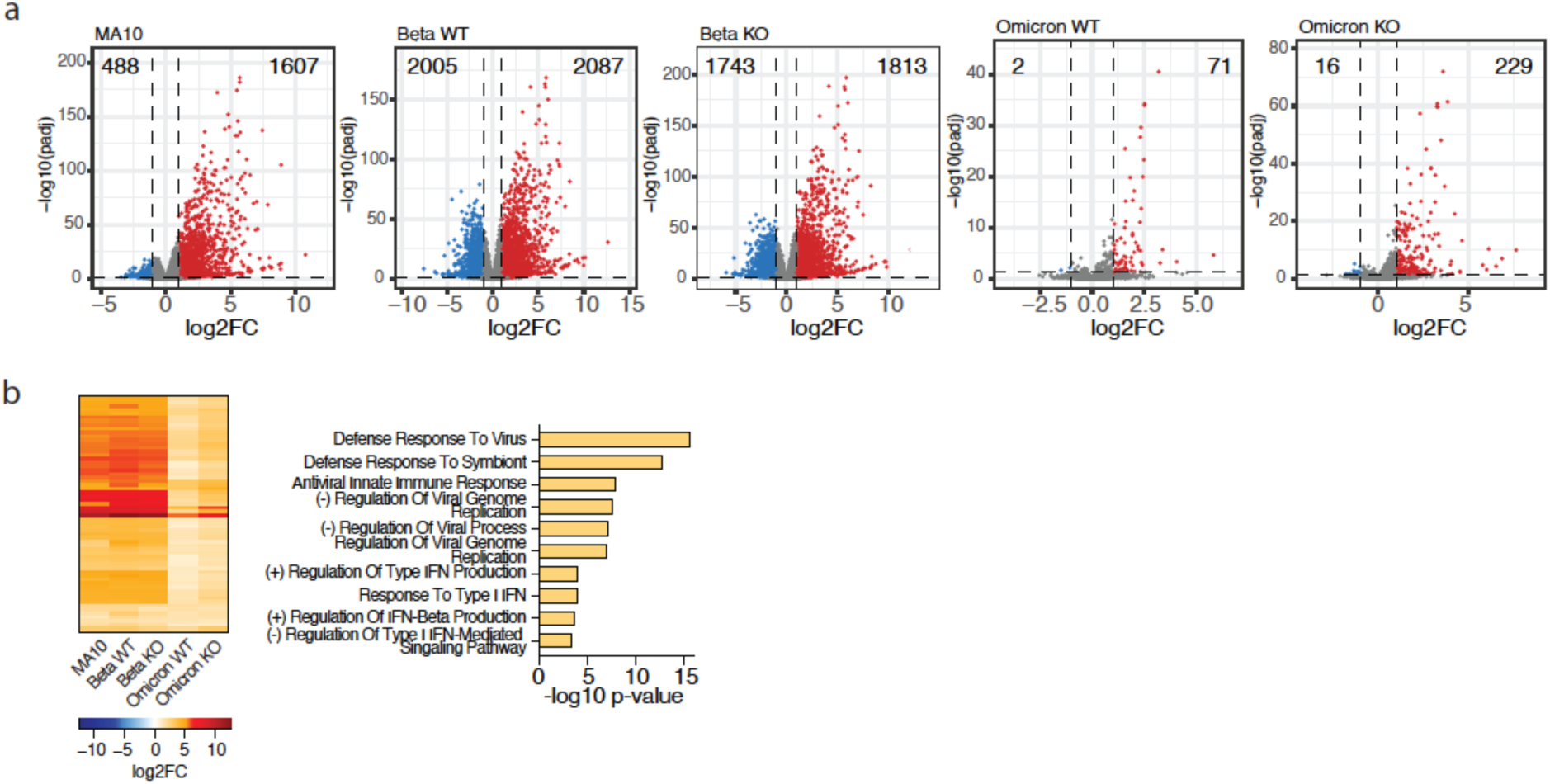
SARS-CoV-2 variants elicit innate immune activation in the lungs of infected mice. (**a**) Volcano plots depict the magnitude (x axis) and significance (y axis) of change in gene expression at day two post-infection. Each data point represents a unique transcript, where color denotes upregulation (red) or downregulation (blue) in expression relative to mock-inoculated mice. Dashed lines depict significance cutoffs (lfc |1|, padj <0.05) and grey data points depict non-significant genes. The number of DEG in either is highlighted in the top corners of the graphs. (**b**) Euclidean distance-based hierarchical clustering of the 63 DEG that overlap across all infectious conditions. Color indicates upregulation (red) in gene expression (left). Bar graphs represent the enrichment scores of GO Biological Pathways terms identified by EnrichR. Each node represents a unique term, edges represent connectivity across terms. Color indicates significant enrichment across infectious groups as indicated by the color scale.

**Supplementary Figure 9.**
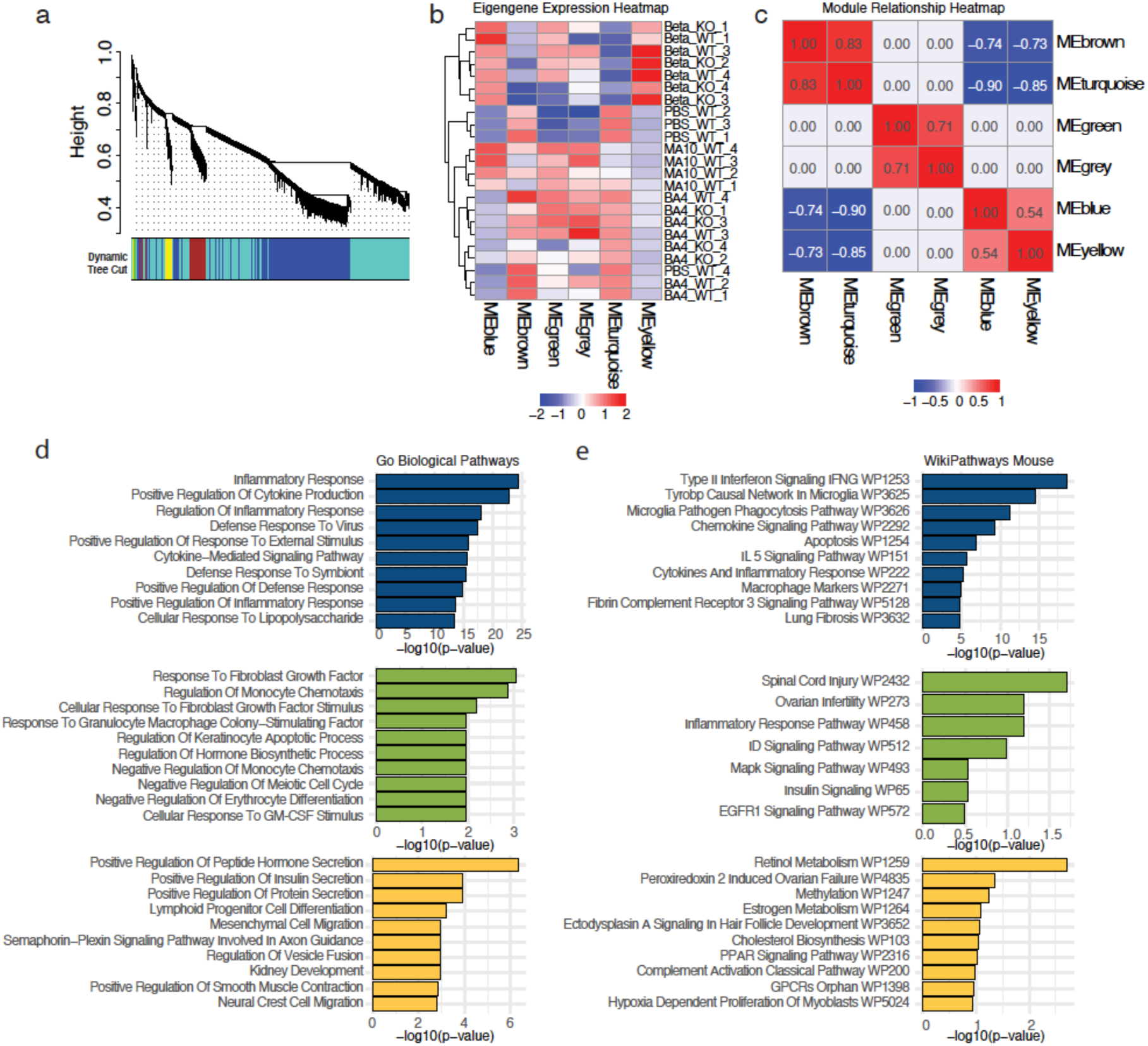
Network analysis reveals gene expression modules associated with viral disease. (**a**) Hierarchical clustering dendrogram of genes with module assignments based on co-expression patterns. Modules were defined using the dynamic tree cut algorithm, and unassigned genes were assigned to the ‘grey’ module. Modules were distinguished by unique colors shown below the dendrogram. (**b**) Heatmap of scaled (Z-score transformation) module eigengene (ME) expression levels with rows representing individual samples, and columns corresponding co-expression network modules. Row-based hierarchical clustering based on their ME expression profiles was performed. The color scale indicates low (blue) to high (red) expression levels. (**c**) Heatmap of significant pairwise Person’s correlations between module eigengenes (MEs). Rows and columns represent modules, clustered based on similarity in their eigengene expression profiles. Non-significant correlations (p > 0.05) were set to zero. The color scale indicates positive (red), negative (blue), or no correlations (white). (**d, e**) Bargraphs represent the -log10 padj significance values of the top (**d**) GO Biological Pathways and (**e**) WikiPathways Mouse terms identified in the ‘brown’ module using EnrichR.

**Supplementary Figure 10.**
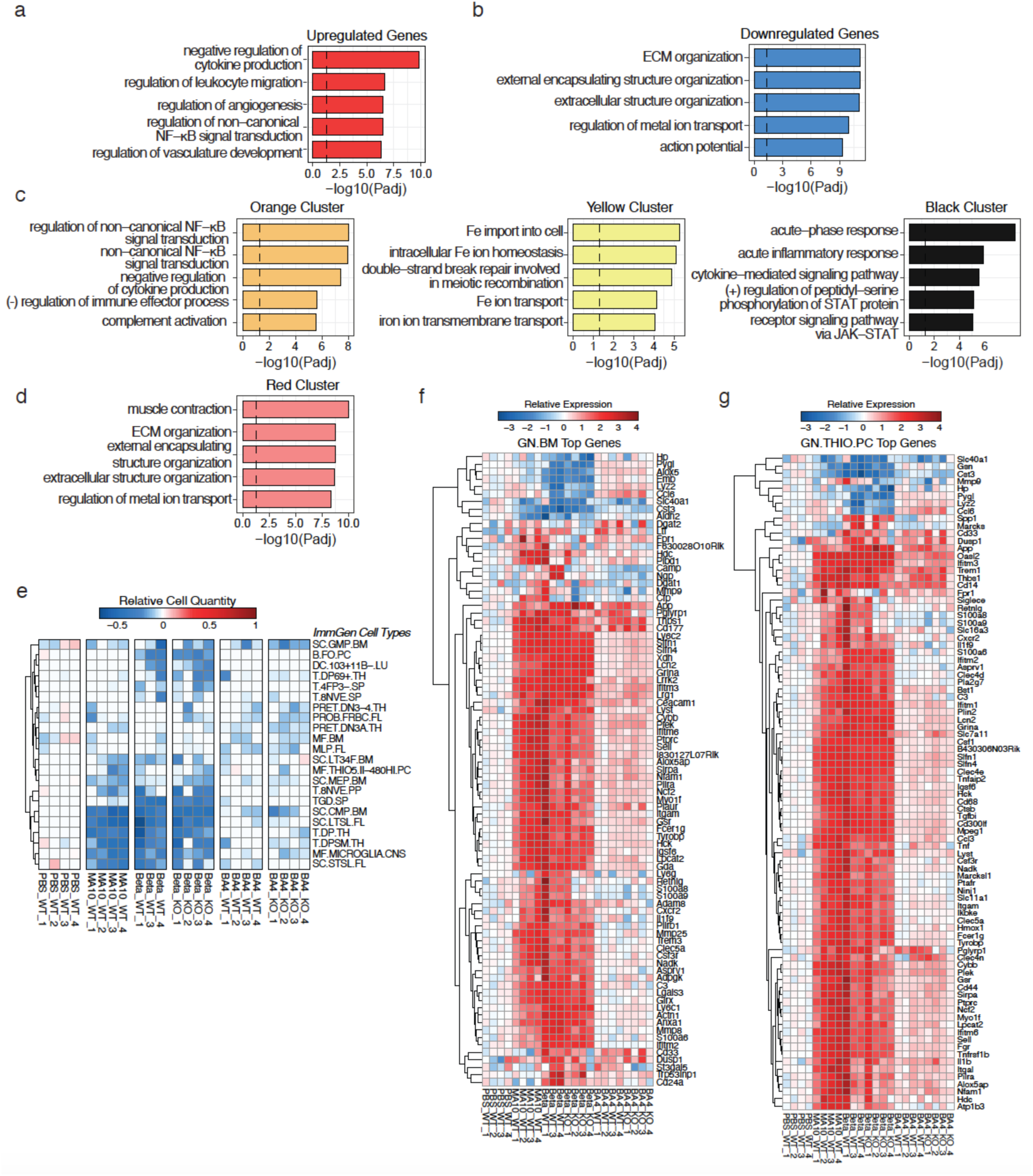
Turquoise model reveals changes in inflammatory responses and extracellular matrix remodeling. Functional enrichment analysis of genes within the Turquiose module. (**a-d**) Bar graphs representing the gene ontology analysis of genes corresponding to (**a**) upregulated, (**b**) downregulated, (**c**) orange, yellow and black, and (**d**) red gene clusters within the Turquoise module identified by hierarchichal clustering in Figure 3A. The x-axis denoted the -log10 padj value of enrichment of the top 10 GO Biological Pathways for each cluster as indicated by color. (**e**) Negatively enriched ImmGen cell types as inferred by DCQ (same as Figure 7f). (**f-g**) Heatmap representing expression, relative to PBS control mice, of the top genes contributing to the (**f**) GN.BM and (**g**) GN.THIO.PC ImmGen cell types in the DCQ algorithm. Red indicates greater expression, and blue indicates lower expression, compared to PBS-inoculated control mice.

